# NFTsim: Theory and Simulation of Multiscale Neural Field Dynamics

**DOI:** 10.1101/237032

**Authors:** P. Sanz-Leon, P. A. Robinson, S. A. Knock, P. M. Drysdale, R. G. Abeysuriya, P. K. Fung, C. J. Rennie, X. Zhao

## Abstract

A user ready, portable, documented software package, *NFTsim*, is presented to facilitate numerical simulations of a wide range of brain systems using continuum neural field modeling. *NFTsim* enables users to simulate key aspects of brain activity at multiple scales. At the microscopic scale, it incorporates characteristics of local interactions between cells, neurotransmitter effects, synaptodendritic delays and feedbacks. At the mesoscopic scale, it incorporates information about medium to large scale axonal ranges of fibers, which are essential to model dissipative wave transmission and to produce synchronous oscillations and associated cross-correlation patterns as observed in local field potential recordings of active tissue. At the scale of the whole brain, *NFTsim* allows for the inclusion of long range pathways, such as thalamocortical projections, when generating macroscopic activity fields. The multiscale nature of the neural activity produced by *NFTsim* has the potential to enable the modeling of resulting quantities measurable via various neuroimaging techniques. In this work, we give a comprehensive description of the design and implementation of the software. Due to its modularity and flexibility, *NFTsim* enables the systematic study of an unlimited number of neural systems with multiple neural populations under a unified framework and allows for direct comparison with analytic and experimental predictions. The code is written in C++ and bundled with Matlab routines for a rapid quantitative analysis and visualization of the outputs. The output of *NFTsim* is stored in plain text file enabling users to select from a broad range of tools for offline analysis. This software enables a wide and convenient use of powerful physiologically-based neural field approaches to brain modeling. *NFTsim* is distributed under the Apache 2.0 license.

## Introduction

The brain is a multiscale physical system, with structures ranging from the size of ion channels to the whole brain, and timescales running from sub-millisecond to multi-year durations. When modeling brain structure and dynamics, it is thus necessary to choose microscale models of individual neurons and their substructures, through network-level models of discrete neurons, to population-level mesoscale and macroscale neural mass and neural field models that average over microstructure and apply from local brain areas up to the whole brain. Many useful results can be obtained analytically from models at various scales, either generally or when applied to specific brain systems and phenomena. However, in order to minimize approximations and make realistic predictions in complex situations, numerical simulations are usually necessary. The purpose of this paper is to present a neural field software package, *NFTsim*, that can simulate scales from a few tenths of a millimeter and a few milliseconds upward, thereby making contact with experiments [1–6] and other classes of simulations over this range [7, 8].

No one type of brain model is optimal at all scales. For example, single neuron models abound in neuroscience, and can include a large number of biophysical effects with relatively few approximations. Many such models have also been used to study networks of interconnected neurons with varying degrees of idealization, thereby revealing a huge number of insights [9, 11, 12]. However, several key problems arise as network size grows: (i) the computational resources required become prohibitive, meaning that simulations can often only be carried out in physiologically unrealistic scenarios, typically with idealized neurons, which may be quantitatively and/or qualitatively inappropriate for the real brain; (ii) it is increasingly difficult to measure and assign biophysical parameters to the individual neurons — e.g., individual connectivities, synaptic strengths, or morphological features, so large groups of neurons are typically assigned identical parameters, thereby partly removing the specificity of such simulations; (iii) analysis and interpretation of results, such as large collections of timeseries of individual soma voltages, becomes increasingly difficult and demanding on storage and postprocessing; (iv) emergence of collective network-level phenomena can be difficult to recognize; (v) the scales of these simulations are well suited to relate to single-neuron measurements, and microscopic pieces of brain tissue, but are distant from those of noninvasive imaging modalities such as functional magnetic resonance imaging (fMRI), electroencephalography (EEG), and magnetoencephalography (MEG) [13–15], which detect signals that result from the aggregate activity of large numbers of neurons; and (vi), inputs from other parts of the brain are neglected, meaning that such models tend to represent isolated pieces of neural tissue.

At the level of neurons and neuronal networks [16, 17], software is abundant, including BRIAN, NEURON, GENESIS, and NeoCortical Simulator [18–23]. A detailed review of tools and implementation strategies for spiking neural network simulations can be found in [9].

At the largest scales, neural mass models average the properties of huge numbers of neurons into those of a single population, without taking account of its spatial aspects. This enables the temporal dynamics of whole neural populations to be approximated, but information on individual neurons and spatial dynamics and patterns is not tracked. This scale can be used to study whole-brain phenomena such as generalized seizures, if time delays within each mass can be neglected. This approach has been used to treat relatively coarse-grained networks of interacting cortical brain regions, each modeled as a neural mass. However, it is rare to see careful attention paid to the need for these representations to approach the continuum limit, in which the cortex is treated as a continuous sheet of neural tissue, as the size of the regions decreases [10, 11, 24, 25], thereby throwing some such discretizations into question. Of course, neural structure is not truly continuous, but its granularity is at a far finer scale than that of the discretizations just mentioned. Above the single-neuron scale and extending to encompass the neural-mass limit as a which properties such as firing rate and soma voltage are viewed as local averages over many neurons, and can vary from point to point, and as functions of time; when correctly discretized, neural mass models are a limiting case of the more general neural fields and should not be viewed as a separate category. Neural fields approximate rate-based models of single neurons from the small scale, while retaining relative timings between neural inputs and outputs. Simultaneously, they self-consistently add spatial structure that is neglected in neural mass models. Hybrid models with features of both neural fields and spiking neurons have also been developed and used to clarify the relationship between these approaches [3], or to enable single-neuron dynamics to be influenced by average neural fields [26], but we do not discuss these classes of models further here.

The issues discussed in the preceding paragraphs are closely analogous to ones that arise in other branches of physics. Specifically, no single model can cover all scales at once. Rather, a hierarchy of models is needed, from the microscale to the macroscale, each relating predictions to measurements at its operational scale. This yields tractable models that can be interpreted in terms of concepts and measurements that apply at the appropriate scales for a given phenomenon. Importantly, each model needs to be related to the ones at nearby scales, especially by making complementary predictions at overlapping scales of common applicability. By analogy, molecular dynamics approaches and statistical mechanics (akin to single neuron approaches) are widely used to track molecules at the microscopic scale, but large-scale theories like thermodynamics and fluid mechanics (akin to neural mass and neural field methods) are more useful and tractable for macroscopic phenomena, and their predictions can be more easily interpreted. At intermediate scales, nonequilibrium thermodynamics and fluctuation theory meet with statistical mechanics and molecular approaches to make complementary predictions of the same phenomena; so that consistency of the various approaches in their common domain can be established. Although molecular-level and spiking-neuron approaches are more fundamental, they are not practical at large scales, and yield results that have to be reinterpreted in terms of larger-scale observables in any case. Conversely, thermodynamic and neural-field approaches fail at spatial and temporal scales that are too short to justify the relevant averaging over a system’s microscopic constituents.

Neural field theory (NFT) incorporates multiple scales such as neurotransmitter effects, synaptodendritic dynamics at the microscale; an average of medium to long-range corticocortical axonal ranges which are essential to model dissipative wave transmission and to produce synchronous oscillations at the mesoscopic scale; and, long-range time delays at the macroscopic scale of the whole brain. Thus, NFT both provides useful macroscopic predictions and can reach down to mesoscopic scales that now overlap with those that can be simulated with neuron-level methods. This provides a range of common applicability on scales of around 1 mm, or slightly less, where complementary predictions can be made and tested – an overlap that will increase as microscopic simulations increase in scale. Equally significantly, quantitative neural field predictions can readily be made of quantities observable by EEG, MEG, fMRI, electrocorticography (ECoG), and other imaging technologies, by adding the biophysics of these signals, measurement procedures, and postprocessing [27–30]. This enables predictions of a single brain model to be tested against multiple phenomena in order to better determine the relevant physiological parameters.

As an illustration of the versatility of NFT approaches, we note that the particular NFT on which the present *NFTsim* software is based has been extensively applied and quantitatively tested against experiments, including EEG, evoked response potentials (ERPs), ECoG, age-related changes to the physiology of the brain, sleep and arousal dynamics, seizures, Parkinson's disease, and other disorders, transcranial magnetic stimulation (TMS), synaptic plasticity phenomena [1, 6,27–39]. Indeed, one of the major strengths of this NFT is its versatility: within the same framework we can express different models to study purely cortical phenomena, the corticothalamic system, basal ganglia, sleep dynamics, or the visual cortex, among an essentially unlimited number of other applications [1, 27–29, 31, 33, 35–38, 40–43]. This NFT has also been clearly linked to hybrid spiking-field approaches [3, 26], and to network and connection-matrix representations of spatial structure in the brain [45], usually obtained via fMRI.

We stress that the NFT embodied in *NFTsim* is not the only possibility. Other NFTs have been developed and applied by numerous authors [46–54], each of which has been applied to one or more physical situations in these and subsequent publications. This list is not exhaustive, since the present work is not intended as a review, but more examples can be found in [11], [25], and [55]. Notably, most of these NFTs can be expressed in the notation of the present paper, and can thus be simulated with the *NFTsim* software described below. Some of these previous neural field models leave out physical effects that are included in *NFTsim*, while others include additional features that remain to be incorporated in a future version of the code.

A few software packages are available to model neural masses and neural fields: [7] developed a neuroinformatics platform for large-scale brain modeling in terms of a network of linked neural masses with anatomically specified cortical geometry [55], long-range connectivity, and local short-range connectivity that approximates the continuum limit when it is Gaussian and homogeneous [24]. While the mathematical framework described in [55] allows for neural field models to be treated using realistic geometry on nonregular grids, a user-ready implementation is not currently available. Similarly, the Brain Dynamics Toolbox [56] provides tools for network-based and continuum models of brain dynamics. The most recent simulation tool for spatiotemporal neural dynamics is the Neural Field Simulator [8], which allows for study of a range of 2D neural field models on a square grid. However, this software does not allow for either the simulation of neural field models with heterogeneous parameters or with multiple populations.

To address the need for research-ready NFT simulation tools with direct application to the study of large-scale brain phenomena, this paper introduces and describes *NFTsim*, a software package that solves neural field equations expressed in differential form for simulating spatially extended systems containing arbitrary numbers of neural populations. The examples of dynamics provided in this work represent perturbations around a fixed point to follow what has been done in previous analytic work. However, *NFTsim* is not limited to the simulation of such dynamics and can produce a range of oscillatory [6, 57], chaotic [58] and bursting dynamics [71].

## Neural Field Theory

Neural field theory (NFT) treats multiscale brain activity by averaging neural quantities such as firing rate, soma voltage, and incoming and outgoing activity over multiple neurons. The scales over which neural field models average must be sufficient to represent large numbers of neurons and spikes, but can still be small enough to resolve quite fine structure in the brain and its activity. *NFTsim* allows an arbitrary number *p* of spatially extended populations of neurons to be simulated. Each of these can be distinguished by its location (e.g., belonging to the cortex or a particular nucleus) and its neural type (e.g., pyramidal excitatory, interneuron). To model a particular system, we must specify the neural populations and the connections between them, including self-connections within a population. If we introduce position and time coordinates **r** and *t*, the main macroscopic variables that describe the activity of neural populations *a* and their interaction with other populations b are: the incoming, axonal spike-rate fields *ϕ_ab_*(**r**, *t*) arriving at population *a* at (**r**, *t*) from population *b*, the dendritic potentials *V_ab_*(**r**, *t*), the mean soma potential *V_a_*(**r**, *t*), the mean firing rate *Q_a_*(**r**, *t*), and the axonal fields *ϕ_ca_*(**r**, *t*) propagating to other populations *c* from population *a*. Figure 1 illustrates the interactions of these quantities: (i) synaptodendritic dynamics involving the incoming axonal fields *ϕ_ab_*(**r**, *t*) to yield the potentials *V_ab_*(**r**, *t*); (ii) dendritic summation and soma charging processes to yield the soma potential *V_a_*(**r**, *t*); (iii) generation of pulses *Q_a_*(**r**, *t*) at the axonal hillock, and (iv) axonal propagation of pulses *ϕ_ca_*(**r**, *t*) within and between neural populations [1]. The following subsections present a review of the equations describing these physiological processes, while Table 1 summarizes the quantities and symbols used in NFT and their SI units. Note that neural field models can be expressed in integrodifferential form [54]. However, in that form, there is always a convolution that is either difficult to handle analytically or numerically [8, 62]. For that reason, all the equations in this NFT [2] and *NFTsim* are expressed and implemented in differential form, respectively.

**Fig 1.**
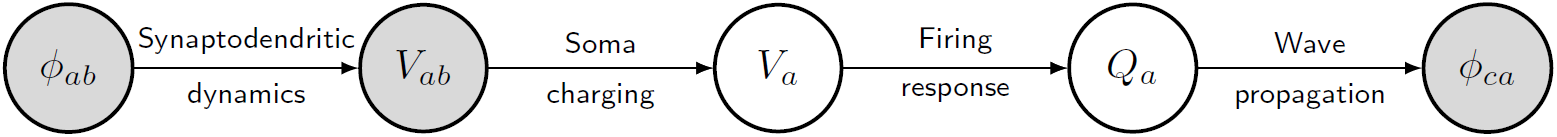
Schematic of the dynamical processes that occur within and between neural populations. Gray circles are quantities associated with interactions between populations (i.e., *a* and *b*), while white circles are quantities associated with a population (i.e., *a* or *b*). Spike-rate fields *ϕ_ab_* arriving at neurons of type *a* from ones of type *b* are modulated by the synaptic dynamics, and undergo dendritic dynamics to produce postsynaptic subpotentials *V_ab_*. These contributions are linearly summed in the dendritic tree, eventually resulting in charging currents at the soma that give rise to the soma potential *V_a_*, after allowing for capacitive effects and leakage. Action potentials generated at the axonal hillock are averaged over a population of neurons. Then, when the mean soma voltage exceeds a threshold, the mean firing rate *Q_a_* of the population is obtained via a nonlinear response function. Finally, the pulses propagate away across the axonal tree and the dendrites of the receiving population *c* as the set of average spike-rate fields *ϕ_ca_*. Note that self-connections with *b* = *a* or *c* = *a* are included.

### Synaptodendritic Dynamics and the Soma Potential

When spikes arrive at synapses on neurons of population *a* from a neural population *b*, they initiate neurotransmitter release and consequent synaptic dynamics, like transmembrane potential changes, followed by dendritic propagation of currents that result in soma charging and consequent modifications of the soma potential. Each of these processes involves its own dynamics and time delays and results in low pass filtering and temporal smoothing of the original spike until the soma response is spread over a time interval that is typically tens of ms, exhibiting a fast rise and an approximately exponential decay [3, 59].

If the overall synaptodendritic and soma responses are linear, which is the most common approximation in the literature [2, 31, 60], the total soma potential *V_a_* is the sum of subpotential contributions *V_ab_*, which are components of perturbation to the dendritic transmembrane potential, arriving at each type of dendritic synapse *ab*. The subscript *a* denotes the receiving population and *b* denotes the neural population from which the incoming spikes originate, distinguished by its source and the neurotransmitter type. The subpotentials *V_ab_* at a particular location comprise contributions from both the wave fields *ϕ_ab_* from other internal populations b and inputs *ϕ_ax_* from external populations *x* [61]; the external inputs are often split into a uniform mean nonspecific excitation and a specific excitation due to structured stimuli. Thus we write the total mean cell body potential as the sum of postsynaptic subpotentials

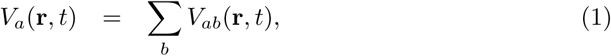
where the subscript *b* distinguishes the different combinations of afferent neural type and synaptic receptor and all the potentials are measured relative to resting [2].

The overall effect of synaptodendritic dynamics and soma charging in response to an incoming weighted pulse-rate field *ϕ_ab_* are well described by an impulse response kernel *L_ab_*(*t* − *t*′)

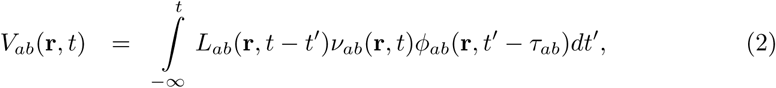

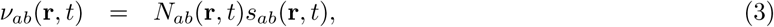
where *ϕ_ab_* is the average rate of spikes arriving at *a* from population *b*; the time delay *τ_ab_* is nonzero when *a* and *b* are in anatomical structures that are separated by a nonzero distance [2]. In Eq. (3), *N_ab_* is the mean number of connections of mean time-integrated synaptic strength *s_ab_* to a cell of type *a* from cells of type *b*. In [2], *L_ab_* is a nonnegative response kernel with

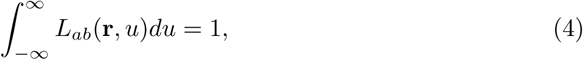
and *L_ab_*(**r**, *u*)=0 for *u* < 0 to express causality. Note that *τ_ab_* are not the only time delays in the system. Propagation delays within a single structure, such as the cortex, are handled by accounting for axonal propagation, as described in section *Propagation of Axonal Pulse-rate Fields*. In *NFTsim L_ab_*(**r**, *t*) is defined as

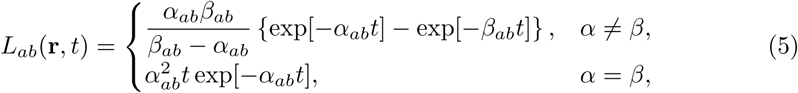
for *t* ≥ 0, with *L_ab_*(**r**, *t*)=0 for *t* < 0 and the **r**-dependence of the positive constants *α* and *β* has been omitted for compactness. These quantities parametrize the decay rate and rise rate of the soma response, respectively, and *β* ≥ *α* is assumed without loss of generality. The temporal profile of the dendritic response function is illustrated in Fig. 2. This function peaks at *t* = ln(*β/α*)/(*β* − *α*) for *α* ≠ *β*; if *α* = *β*, the peak is at *t* =1*/α*. In addition, there are two special cases of Eq. (5): (i) if either *α* →∞ or *β* →∞, then *L_ab_* becomes a single exponential function in which only one of the characteristic timescales dominates; and, (ii) if *α* = *β* = ∞, then the kernel reduces to the impulse *L_ab_*(**r**, *t*)= *δ*(**r**, *t*).

**Fig 2.**
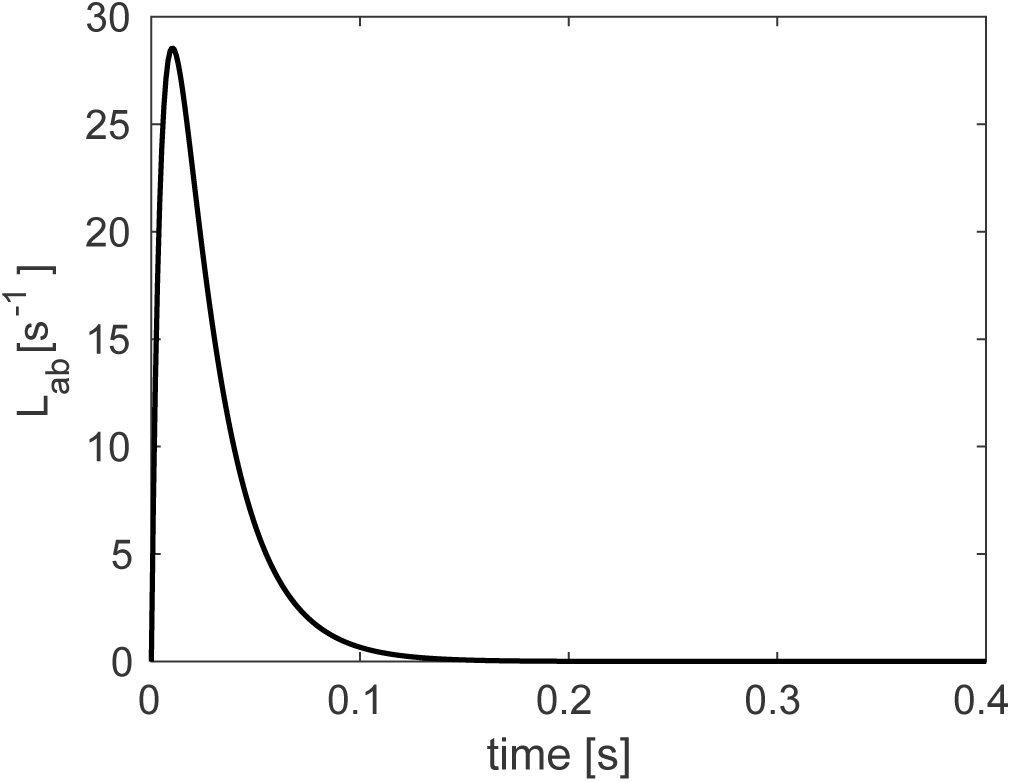
Dendritic response function. The response to a delta-function input, via *L_ab_* as defined in Eq. (5), for decay rate parameter *α_ab_* =45 s^−1^ and rise rate parameter *β_ab_* = 185 s^−1^. This function peaks at *t* = ln(*β/α*)/(*β* − *α*) for *α* ≠ *β*.

The convolution in Eq. (2) can be re-expressed as

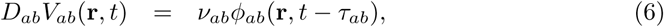
where the differential operator *D_ab_* is given by

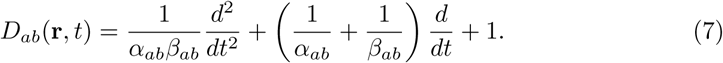

In some previous work [59] a special approximation has been used where *α_ab_* and *β_ab_* are independent of *b* and are thus treated as effective values, representing an average over different receptor time constants. Under this approximation Eq. (6) becomes

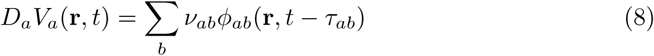

All the aforementioned cases and forms of the operators (differential and integral) are implemented in *NFTsim*.

### Generation of Pulses

Pulses (i.e., spikes or action potentials) are produced at the axonal hillock when the soma potential exceeds a threshold potential *θ_a_*(**r**, *t*). When we consider the mean response of a population of neurons to a mean soma potential we must bear in mind that each neuron has slightly different morphology and environment. Hence, they respond slightly differently in the same mean environment. This has the effect of blurring the firing threshold and the resulting overall population response function is widely approximated by the nonlinear form [49]

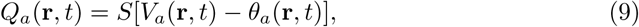
where *θ_a_* is the mean threshold potential of population *a* and *S_a_* is a function that increases monotonically from zero at large negative *V_a_* to a maximum firing rate 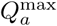 at large positive *V_a_*, with the steepest increase concentrated around the mean threshold *θ_a_*. *NFTsim* employs by default the nonlinear sigmoid response function

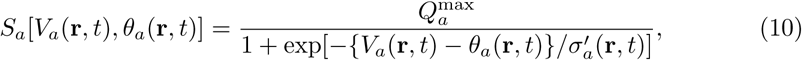
where 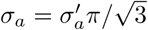 is the population standard deviation of the soma voltage relative to the threshold. If the function in Eq. (10) is linearized to consider small perturbations around a steady state of the system [2, 32], one finds the linear response function

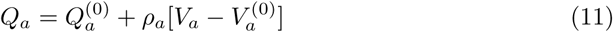
where 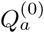 and 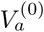 are the relevant steady-state values and *ρ_a_* = *dQ_a_/dV_a_*, is the slope (0) of the sigmoid function, evaluated at 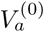 [2]. This linear population response function is also implemented in *NFTsim* and other functional forms can be defined as well.

### Propagation of Axonal Pulse-rate Fields

The propagation of the pulses *Q_b_*(**r**, *t*) in each population *b* generates an outgoing mean field *ϕ_ab_* that propagates via axons to the population *a* at other locations. In general, this propagation can depend on both the initial and final populations, and can incorporate arbitrary nonuniformities and a range of propagation velocities via propagator methods, for example [62, 63]. However, considerable theoretical and experimental work has shown that, to a good approximation, the mean field of axonal signals in a smoothly structured neural population propagates approximately as if governed by an isotropic damped wave equation [2, 48, 50, 53, 54, 64–69]. In *NFTsim* we implement the widely used equation

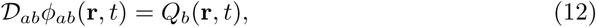
with

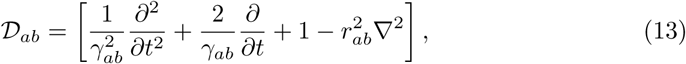
where *γ_ab_* = *v_ab_/r_ab_* is a temporal damping coefficient, *r_ab_* is the spatial effective axonal range, *v_ab_* is the axonal velocity [2, 54, 65–69], and ∇^2^ is the Laplacian operator. Equations (12) and (13) constitute the two-dimensional generalization of the telegrapher’s equation [2, 54, 70]. More generally, *γ_ab_*, *r_ab_*, and *v_ab_* can be functions of position. If the special case of spatially uniform activity is considered, the Laplacian operator has no effect and can be omitted from (13). This special case results in the harmonic operator

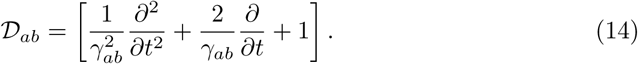

We stress that this is not the same as using a local neural mass model because the damping parameter *γ_ab_* depends on spatial propagation. To obtain the neural mass limit, one also needs to set the spatial ranges *r_ab_* =0 so *γ_ab_* becomes infinite and

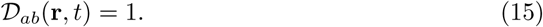

This yields

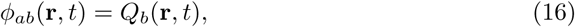
which is termed the *local interaction approximation* [2, 51].

The parameter *r_ab_* in the propagators in Eqs (13) and (14) encompasses divergence of axons traveling to the target population *a* from the source population *b* and the extent of dendritic arborization of the target population *a*, and thus *r_ab_* ≠ *r_ba_* in general [82].

## Design and Implementation of NFT*sim*

This section presents a comprehensive description of *NFTsim*. The subsection *General Workflow* gives an overview of the typical usage workflow of *NFTsim*. The subsection *Classes and their Interactions describes the main NFTsim* classes, which represent the biophysical processes and quantities introduced in *Neural Field Theory*.

**Table 1.** NFT quantities and associated SI units. Symbols used in NFT, associated physical quantities and their SI units. Double subscripts ab mean that the target population is a and the source population is b.

| Symbol | Description | Units |
| --- | --- | --- |
| $Q_a^{\max}$ | Maximum firing rate | $s^{-1}$ |
| $\gamma_{ab}$ | Damping rate | $s^{-1}$ |
| $v_{ab}$ | Wave velocity | $m\ s^{-1}$ |
| $r_{ab}$ | Mean axonal range | m |
| $\theta_a$ | Mean neural firing threshold | V |
| $\sigma'_a$ | Standard deviation of the firing threshold | V |
| $\alpha_{ab}$ | Mean dendritic response decay rate | $s^{-1}$ |
| $\beta_{ab}$ | Mean dendritic response rise rate | $s^{-1}$ |
| $\nu_{ab}$ | Synaptic coupling strength | V s |
| $\tau_{ab}$ | Long range time delay | s |
| $\phi_{ab}$ | Axonal field | $s^{-1}$ |
| $Q_a$ | Mean firing rate | $s^{-1}$ |
| $V_{ab}$ | Subpotential | V |
| $V_a$ | Mean soma potential | V |

Next, subsection *Input-Output illustrates with examples how to specify a model in the* input configuration file to *NFTsim* and how to interpret the output file. In addition, subsection *Numerical Methods, Considerations, and Constraints elaborates on the* numerical approaches and constraints used to correctly solve the equations of neural field models while attaining numerical accuracy and stability. Table 2 summarizes the configuration parameters relevant to these methods. Lastly, subsection *Analysis and Visualization* presents a simple example of how to run a simulation, and analyze and visualize the results using the auxiliary Matlab module `+nf`. A list of the available functions in this module is presented in Table 3.

The typographic conventions used in the remainder of this text are that: (i) all computer code is typeset in a typewriter font; and (ii) code snippets are framed by horizontal lines with line numbers on the left.

### General Workflow

A typical *NFTsim* workflow consists of three broad phases: configuration; simulation; and postprocessing. The first phase involves writing a configuration file that specifies the neural field model as well as other parameters required to run a simulation. This file is a human readable plain text file with the extension `.conf`. Once a configuration is specified the simulation can be launched by invoking the `nftsim` executable, either directly via a shell (eg. *bash*) terminal

~~~
user@host $ nftsim -i <my-model.conf> -o <my-model.output>
~~~

or indirectly via the `nf.run` Matlab function. In the simulation phase, *NFTsim* reads the configuration file, specified after the flag `-i`, builds the objects of the specified model, runs the simulation and writes the output file, which contains the timeseries of the neural quantities requested in the configuration file. The name of the output file can be specified using the flag `-o` and must have the extension `.output`. In the absence of an output file name, *NFTsim* uses the input file name with the extension `.output`. For autogenerated output file names, the flag `-t` can be used to append a string to the YYYY-MM-DDTHHMMSS, which follows the standard ISO 8601 [76] to represent date and time. In the postprocessing phase, the simulation results can be analyzed offline and visualized with the functions provided in the Matlab module `+nf`.

### Code Architecture

Neural field models can be decomposed into a small number of objects, that represent their various parts. Each object has intrinsic properties that, in turn, can be well represented as classes, each of which is a set of elements having common attributes different from other sets, using object oriented programming. *NFTsim* classes have been implemented in C++ (C++11 standard) [77, 78].

The most prominent components of neural field models are populations, synaptic connections, and propagators. Each of these components (or objects) is described by a main base class with properties specific to a group of objects. Derived classes are defined via the mechanism of class inheritance which allows for: (i) the definition of class in terms of another class; (ii) the customization of different parts of the system being modeled; and (iii) the extension of the functionalities of the library. For instance, a base class describing propagators has properties such as axonal range and axonal velocity. These properties are common to different propagators (derived classes) such as the wave propagator in Eq. (13) or the harmonic propagator in Eq. (14), and are inherited from the base class. However, the optimal method to solve each form of propagation may vary and thus each propagator-derived class can have its own solver. Furthermore, there are auxiliary base classes that define additional properties of the main classes described above. These auxiliary classes embody processes like dendritic dynamics, soma charging, firing response, external stimuli, and anatomical time delays.

Thanks to this modular architecture, *NFTsim* allows for the specification of models with (i) an arbitrary number of neural populations, of different types and with different parameter sets; (ii) different types of connections between pairs of populations; and (iii) different types of activity propagation, with or without propagation time delays between and within neural populations.

### Classes and their Interactions

An overview of *NFTsim*’s calling interactions between classes, is illustrated in Fig. 3. In this diagram main and auxiliary base classes are positioned so that, in a simulation, their position corresponds to being initialized and stepped forward in time from top to bottom and from left to right within each row. In the first row, we see the high-level class `Solver` which coordinates how the other classes interact during a simulation. In the second row, the main base class `Propagator` computes each of the axonal pulse-rate fields *ϕ_ab_* generated by the firing rate *Q_b_*. In any given neural field model there are as many `Propagator` objects as there are connections. These can be any of three derived `Propagator` classes (`Wave`, `Harmonic`, `Map`) implemented to accommodate the operators defined in Eqs (13), (14), or (15), respectively. The `Wave` class uses an explicit time stepping method based on second order central difference schemes in space and time (see Explicit Difference Method and Boundary Conditions for the 2D Wave Equation). The `Harmonic` class implements Eq. (14), where for spatially homogeneous models the Laplacian term is zero and one finds a damped oscillator response. This class uses a standard fourth-order Runge-Kutta (RK4) explicit forward time stepping method with a fixed time step [86]. Lastly, the `Map` class, where the propagator is simply a direct mapping as in Eq. (15). Below `Propagator`, there is the auxiliary class `Tau`, which handles the activity history and retrieves the appropriate delayed activity when the discrete time delay *τ_ab_* is nonzero. This class actually stores time delayed activity between every pair of populations, for every spatial location, and makes it available to the numerical solvers. Then, to the right of `Propagator`, the `Coupling` class handles synaptic connections and their dynamics. The base `Coupling` class assumes that the synaptic strengths are constant over space and time. Thus, the output signal is a product of incoming activity and synaptic weights. Other derived `Coupling` classes implement temporally varying synaptic strengths as in [36], or modulation by pre-or postsynaptic activity, as in [40]. To the right of `Coupling`, the `Population` class describes neural population activity and its parameters define the type.

**Fig 3.**
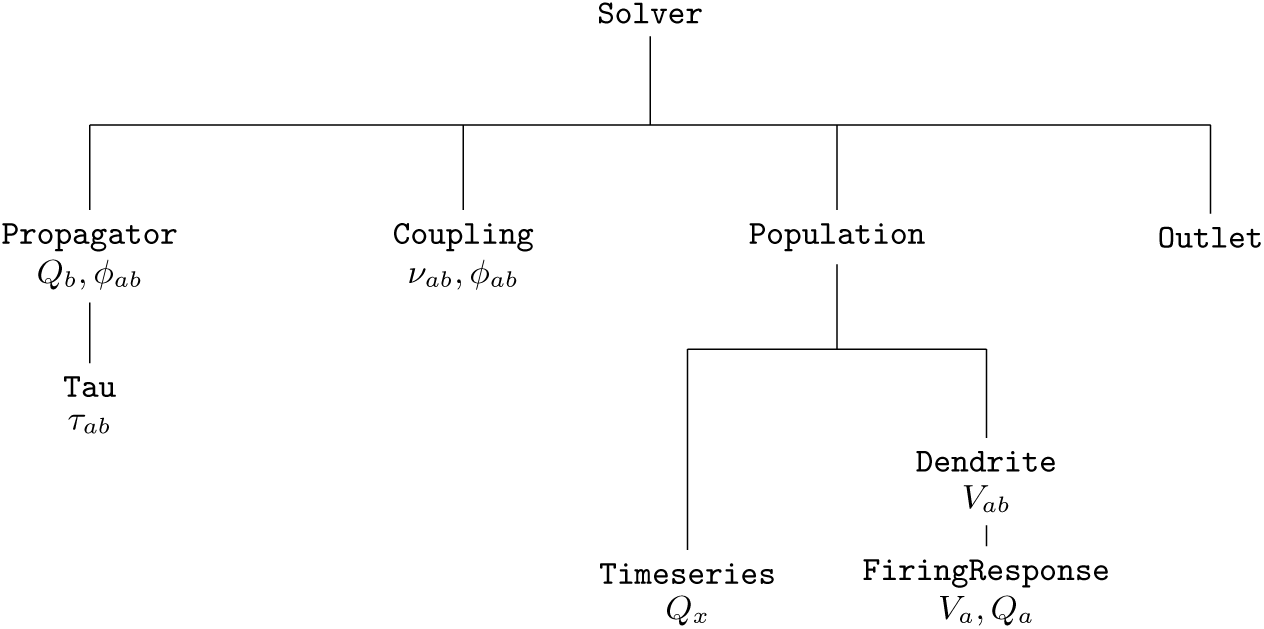
Simplified diagram of *NFTsim’s* call graph. The execution of a simulation is controlled by the class `Solver`. Initial conditions are given in terms of firing rates *Q_b_* which are then propagated to other populations via `Propagator`. Synaptic connections are handled via `Coupling`. The incoming activity to postsynaptic `Population` undergoes dendritic dynamics via `Dendrite`. The sum of individual contributions *V_ab_* and the resulting firing response are handled by `FiringResponse`. The class `Timeseries` is used to represent external inputs *Q_x_* from a stimulus `Population`. Lastly, the class `Outlet` stores the variables that are written to the output file.

In the third and fourth rows, below `Population`, we see that each `Population` uses two subsidiary classes: an array of `Dendrite` objects (one for every population connected via a `Coupling`); and, a `FiringResponse`. The signal from a `Coupling` object is passed to a corresponding `Dendrite` object which implements the synaptodendritic effects defined in Eq. (6). The contributions *V_ab_* are then summed to yield the soma potential *V_a_* of the population. Then, the population’s `FiringResponse` object implements Eq. (9) to calculate the resulting population firing rate *Q_a_*. Different forms of the activation function are specified within the base `FiringResponse` class. Other types of activation function that involve modulation of parameters due to presynaptic or postsynaptic activity are implemented in classes derived from the `FiringResponse` class. Such is the case of `BurstingResponse` that implements modulation of firing threshold *θ_a_* [71]. External or stimulus populations are also objects of the `Population` class. However, their activity is a predefined spatiotemporal profile of firing rate *Q_x_*, that represents a chosen input and is contained in an object of the class `Timeseries`. In *NFTsim* the external inputs may include noisy and/or coherent components which may or may not be spatially localized (e.g., afferent to the visual thalamus in response to a visual stimulus). Currently, *NFTsim* supports a number of different external driving signals (*ϕ_ax_*) to stimulate any population *a* of a system. These signals include: a constant value equivalent to applying DC voltage; sine waves; square pulse trains; and, Gaussian white noise to simulate random perturbations. These basic functions can be combined additively to generate more complex stimulation signals.

**Fig 4.**
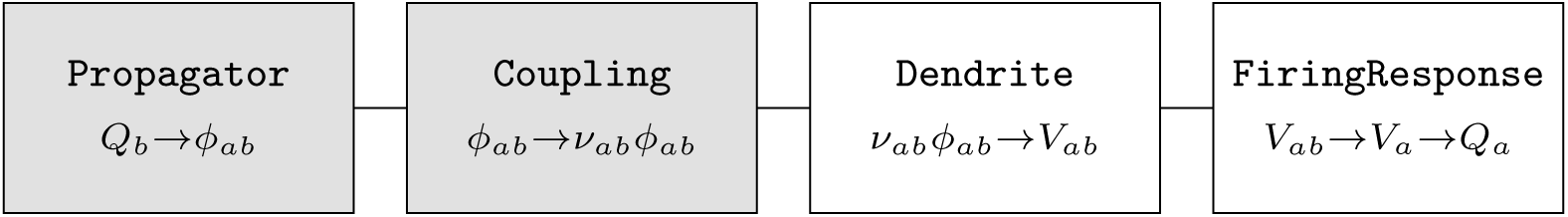
*NFTsim* classes associated with biophysical processes. This diagram illustrates the relationship of the classes in the library and the biophysical transformations they represent. Input variables are on the left, while output variables are on the right. Gray boxes are classes associated with interactions between populations, while white boxes are classes associated with internal mechanisms of a population.

Lastly, to the right of `Population`, the class `Outlet`, stores the variables that are written to the output file.

In summary, a compact representation of the neural field equations with the label of the associated *NFTsim* classes is

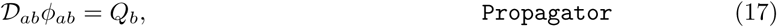

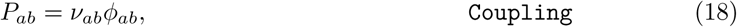

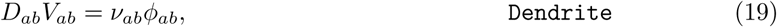

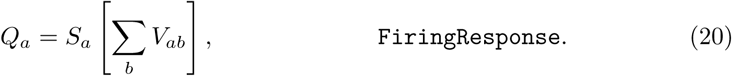
where the auxiliary variable *P_ab_* in Eq. (18) is only defined inside the `Coupling` class and assigned the presynaptic inputs weighted by the local synaptic coupling strength. Figure 4, which is analogous to the diagram presented in Fig. 1, illustrates the input and output variables of each class and the direction in which they flow within a simulation.

### Input-Output

The main routine of *NFTsim* takes a plain text configuration file as input, where all the model description and simulation parameters are specified, and writes the simulation result to an output file. Both the configuration file and output file are plain text files, so launching simulations and reading the results with other programming languages is also possible. Note that all the parameters in the configuration and output files are specified directly in SI units without prefixes (e.g., s, s^−1^, V); e.g., a value of 1 mV is written as `1e-3` (where V is implicit).

#### Configuration and Output Files

The following listing shows an exemplar configuration file, named `e-erps.conf`, which is included with other examples in the `configs/` directory of *NFTsim*. This file specifies a neural field model with a single cortical excitatory population that receives inputs from an external population which is the source of a stimulus to the cortex. In this example, parameters were taken from [33], with the exception of the axonal propagation parameters, which are tuned to emphasize wave propagation properties (i.e., by decreasing the damping rate *γ_ab_*). We emphasize that this an illustrative example, and that while it emulates the scenario of evoked responses due to a stimulus (e.g. a flash of light) it does not represent any specific experiment.

The cortical population is initially in a steady state of low firing rate around 10 s^−1^ and is driven by two pulses applied toward the center of the grid. The first pulse occurs at *t* ≈ 32 ms and has a positive amplitude of 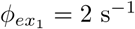. The onset of the second pulse is *t* ≈ 60 ms and has a negative amplitude of 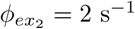.

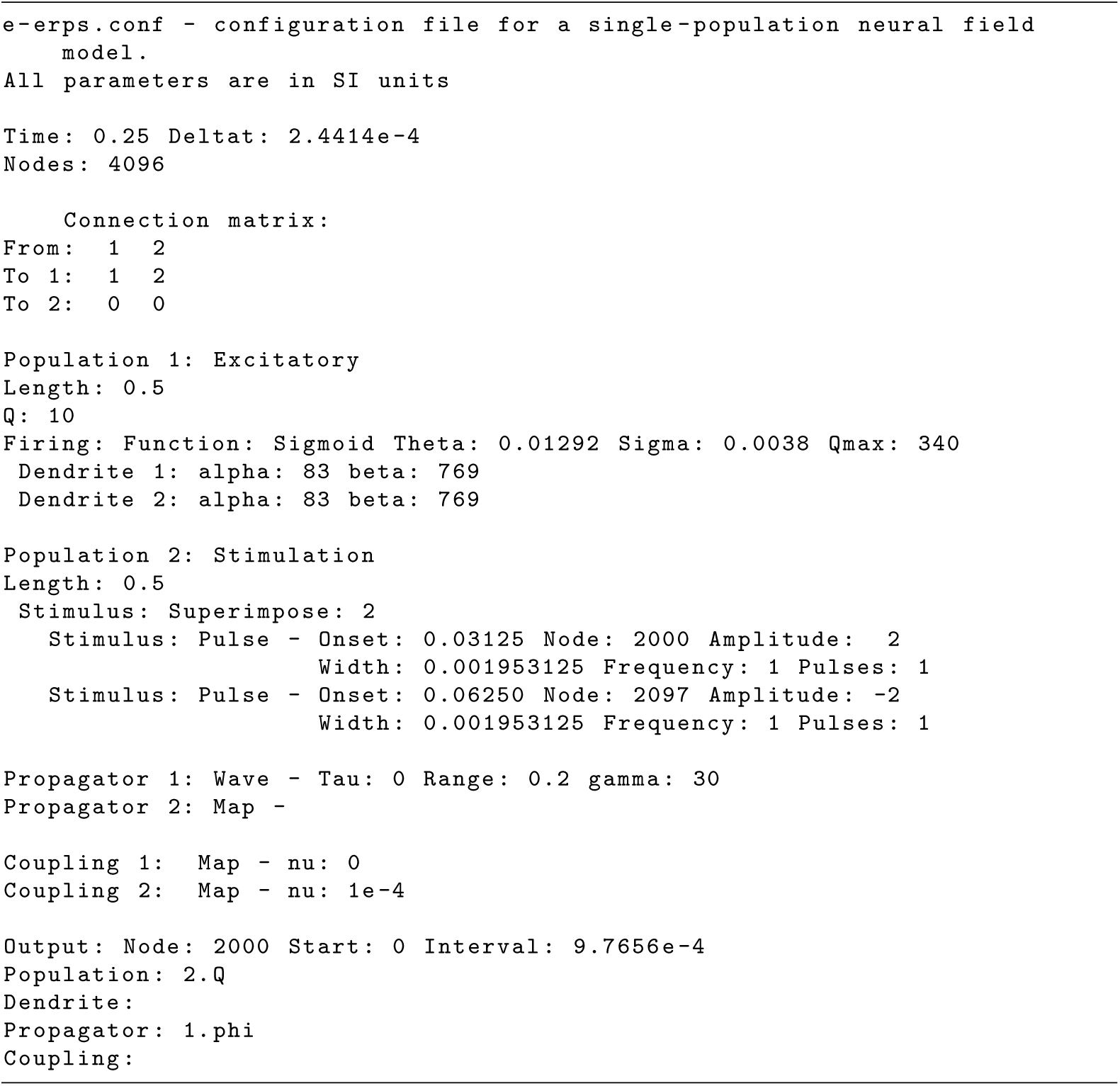

The above file starts with a brief description of the model to be simulated. This comment is optional and can span multiple lines. In lines 4–5, global parameters for the simulation are defined: simulation duration (`Time`), time step size (`Deltat`), and the total number of nodes in the two dimensional grid (`Nodes`).

The aforementioned parameters are followed by the specification of a square connection matrix in lines 7–10, where the rows are the target populations and columns indicate the source populations. In this matrix, a positive integer indicates there is a connection between two populations and it also serves as an identifier of that connection. In the case presented above, there are only two nonzero connections, connection `1` to `Population 1` from itself and connection `2` to `Population 1` from `Population 2`. The couplings, dendrites and propagators are labeled by these consecutive positive integers. The two populations of this example are defined in lines 12–25. Each population in the model is specified separately, indicating its type (e.g., excitatory, inhibitory, or external), the physical size of its longest side (`Length`), its initial condition in terms of firing rate `Q`, and its type of dendritic and firing responses. The next step, in lines 27–28, is to define the type of propagation and coupling between each pair of connected populations. In line 27, the axonal propagation of the excitatory-excitatory connection follows a damped wave equation, with zero long-range time delay (`Tau`), characteristic spatial range of 0.2 m (`Range`) and a damping coefficient of 30 s^−1^ (`gamma`). Finally, at the end of the configuration file, from line 33 onwards in this example, we specify which timeseries are written to the output file.

There are three global output parameters: `Node` which specifies the labels of the grid nodes whose activity will be written to the output file; `Start`, sets the time (in seconds) from which the output timeseries will be written, and cannot be larger than the total simulation duration `Time`; and, `Interval` is the sampling interval between points in the output timeseries. The `Interval` should be chosen as an integer multiple of `Deltat`, that is, the ratio `Interval`/`Deltat` should be an integer number *K*, because *NFTsim* does not perform interpolation or averaging when users request downsampled output. The timeseries written to disk is simply a subsampled version of the original, where only every *K*-th sample is kept. In lines 34 and 36 we see that *NFTsim* has to write the firing rate (`Q`) of `Population 2`, and the axonal field `phi` of `Propagator 1`, respectively. *NFTsim* first writes the configuration file at the top of the output file to ensure full reproducibility of the results, then it writes a line filled with the symbol `=`, and finally, it writes the requested timeseries. Below we show an excerpt of the output file `e-erps.output`.

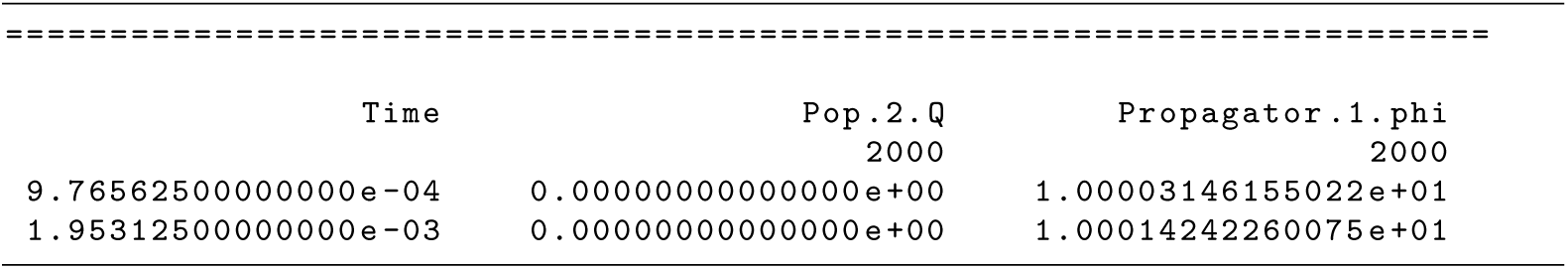

Here, the first column is the time vector. The values are expressed in seconds. The second column is the firing rate `Q` of the second population at node 2000. The third column is the excitatory field of `Propagator 1` from `Population 1` to itself at node 2000. Line 3 provides the label of each timeseries, while line 4 shows the node index.

### Numerical Methods, Considerations, and Constraints

This section focuses on considerations and constraints regarding the numerical methods implemented in *NFTsim*. In *Initial Conditions we give a general overview and strategies* to set initial conditions for neural field simulations. Furthermore, *Discretization of the Spatial Domain* and *Courant Condition describe the way space is discretized in NFTsim* and the maximum grid ratio for correctly solving the 2D damped wave equation, respectively. In *Explicit Difference Method and Boundary Conditions for the 2D Wave Equation* we explain the stepping method used to solve the wave equation on a finite grid. Lastly, in *Time Delays*, we briefly explain how time delays are handled in *NFTsim*.

#### Initial Conditions

Neural field equations are partial delay differential equations (PDDEs), thus at the start of a simulation activity from previous times is required for initialization. *NFTsim* assumes the system is initialized at a stable fixed point and then fills a history array, which stores the past activity of the system, with the values of firing rate at equilibrium. A more detailed explanation on how time delays are handled is given in a subsequent section.

In a steady state all the temporal derivatives can be set to zero. Furthermore, *NFTsim* currently also assumes that the initial activity is uniform spatially, so the spatial derivatives are also set to zero. Under these assumptions, if the initial conditions are not exaclty a stable state of the system, one can expect to see transient activity until the system settles into the closest stable attractor (either a fixed point or another manifold).

In NFT, the number of stable steady-state equilibria strongly depends on the number of populations and connections between them [79], thus providing a general method to find the steady-state solutions is beyond the scope of *NFTsim*’s functionality.

Nevertheless, there are four main strategies that users may adopt to set initial conditions:

i. Finding the steady-state solutions analytically. This method is successful for simple neural field models with a couple to a few populations such as a purely cortical model [32], and that do not include a nonlinearity in their firing response.
ii. Finding steady-state solutions numerically. If one uses the NFT equations which include a nonlinear firing response, then the steady-state equation is a transcendental function of either firing rate or voltage, and its roots cannot be calculated analytically. Fixed points can be identified by evaluating the steady state equation of the system between consecutive test values of one of the fields or voltages e.g., 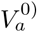 or 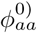), and detecting the zero crossings. This is the method that has been used extensively to find the roots of the corticothalamic model for different parameter ranges [6, 29, 30, 39, 79]. A standard root finding algorithm (e.g., Newton-Raphson) can then be used to refine the roots. If the steady-state equation of the neural field model depends on more than one variable [57] then a root finding algorithm like Broyden’s method is required. Note that the strategy described here does not identify the stability of the fixed points.
iii. Running auxiliary simulations. This approach is best suited for scenarios in which one already has an initial estimate of the initial stable state of the system; and for nonuniform situations [80, 81], in which case the auxiliary simulations are run for the uniform case and the nonuniformities in the parameters are introduced in the main simulations. Auxiliary simulations should be long enough to give the system enough time to reach a stable state. The end state of this auxiliary simulation can then be used to provide the initial conditions for other simulations.
iv. Using Monte Carlo methods to run numerous simulations in *NFTsim* with randomly sampled initial conditions in order to find the stable states. This approach is more general than the previous one and does not require any a priori knowledge of the initial conditions. This approach is best suited for neural field models with several populations and for which finding the steady states of the system following (i) or (ii) is not possible or is too cumbersome. If multiple stable steady states are found [48, 79], users must decide which one is to be used for the main simulations. In NFT, the linearly stable fixed point that represents the lowest firing rates is usually selected as the initial condition on the basis that represents a normal brain state [2, 29].

#### Discretization of the Spatial Domain

Each population is modeled as a 2D rectangular sheet. In *NFTsim*, the physical spatial domain of each population, whatever its extent, is divided into a finite number *N* of uniform grid cells (or nodes), which remain invariant throughout the simulations for all times. Note that in this work grid cells refer to smallest surface area units used to discretize a continuous and spatially extended domain such as the cortex, and not to the homonymous biological cells in the enthorinal cortex.

In a configuration file, the parameter `Length` corresponds to the physical length of the *x*-axis. By default, the domains are assumed to be square with *L_x_* = *L_y_*. In this case, the value of the parameter `Nodes` must be a perfect square so that the spatial resolutions

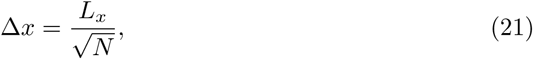
and

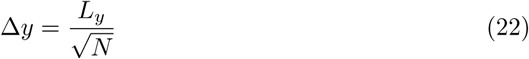
are the same.

To define a rectangular domain, in addition to the parameter `Nodes` (*N*), in the configuration file one can specify the number of nodes along the *x*-axis via the parameter `Longside nodes` (*N_x_*). In this case, the number of nodes along the *x* and *y* axes are different, but the spacing remains the same for both axes (i.e., Δ*x* = Δ*y*)

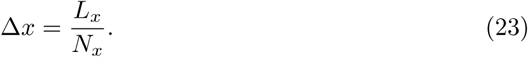

The number of nodes and physical length of the *y*-axis can be obtained as *N_y_* = *N/N_x_* and *L_y_* = *N_y_*Δ*y*, respectively. Table 2 summarizes the symbols and configuration length and size parameters used in this section and in the remainder of the text.

As an example, we show part of a configuration file for a neural field model with two populations. The physical length of the first population 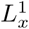 is larger than the length of second population 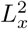.

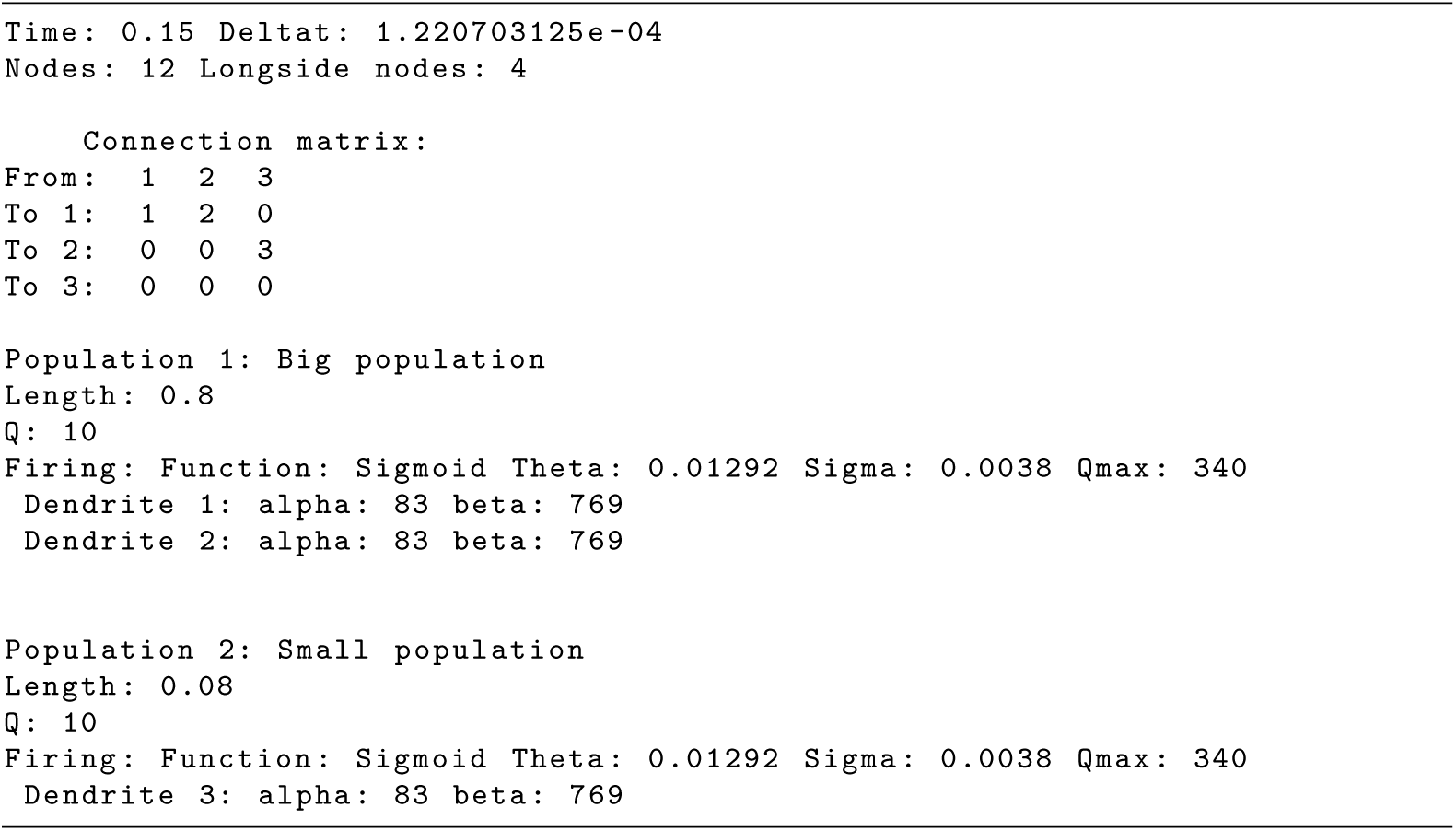

In the above file the two internal populations are modeled as rectangular grids with a total of 12 nodes or grid cells, and with the number of nodes of the longest side specified by `Longside nodes`. The resulting 2D grid has a size of 4 × 3 nodes as shown in the schematic of Fig. 5. For illustrative purposes, the parameter values used in this configuration file have been exaggerated so the link between the input parameters and the discretization of the space shown in the schematic is clear. However, this configuration file will not produce accurate results because the spatial resolution is too coarse.

Figure 5 illustrates that *NFTsim* populations are linked via a primary topographic one-to-one map, which implies that all the populations must have the same number of grid points *N*, even if they have different physical spatial dimensions. We assign the same map coordinate **r***_n_* to homologous grid cells in different populations. In this example, **r**_1_ is assumed to be the actual physical position in `Population 1`, but in `Population 2`, **r**_1_ denotes a rescaled physical dimension. Also, any physical position **r***_n_*, for *n* =1, …, *N* is assumed to be at the center of a grid cell, which is also labeled with integers *n* =1, …, *N*. For instance, in `Population 1`, **r**_1_ corresponds to position (Δ*x*_1_=2, Δ*y*_1_=2) = (0.1, 0.1) m; and, in Population 2, **r**_1_ corresponds to position (Δ*x*_2_/2, Δ*y*_2_/2) = (0.01, 0.01) m.

**Table 2.**
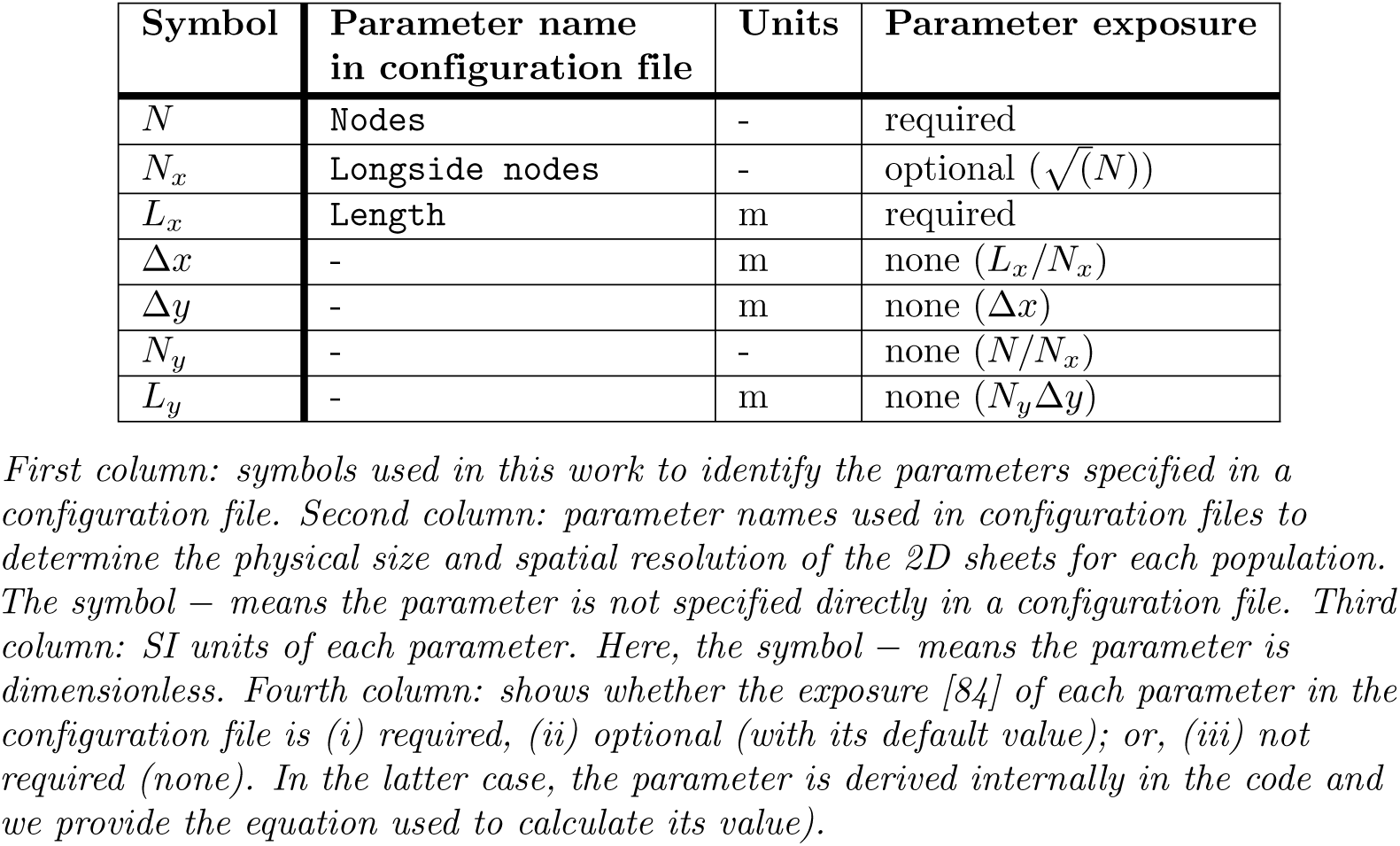
Symbols, configuration parameters and units.

| Symbol | Parameter name in configuration file | Units | Parameter exposure |
| --- | --- | --- | --- |
| $N$ | Nodes | - | required |
| $N_x$ | Longside nodes | - | optional ($\sqrt{(N)}$) |
| $L_x$ | Length | m | required |
| $\Delta x$ | - | m | none ($L_x/N_x$) |
| $\Delta y$ | - | m | none ($\Delta x$) |
| $N_y$ | - | - | none ($N/N_x$) |
| $L_y$ | - | m | none ($N_y \Delta y$) |

First column: symbols used in this work to identify the parameters specified in a configuration file. Second column: parameter names used in configuration files to determine the physical size and spatial resolution of the 2D sheets for each population. The symbol – means the parameter is not specified directly in a configuration file. Third column: SI units of each parameter. Here, the symbol – means the parameter is dimensionless. Fourth column: shows whether the exposure [84] of each parameter in the configuration file is (i) required, (ii) optional (with its default value); or, (iii) not required (none). In the latter case, the parameter is derived internally in the code and we provide the equation used to calculate its value).

#### Courant Condition

The interval Δ*x* is used to evaluate whether the current parameters satisfy the Courant condition, a necessary condition for obtaining stable solutions when solving hyperbolic partial differential equations on a regular discrete grid. For the wave equation in 1D the dimensionless number

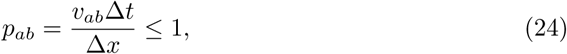
is called the Courant number [85]; Δ*t* is the integration time step size and *v_ab_* = *γ_ab_r_ab_* is the magnitude of the wave velocity. In the continuum wave equation, activity propagates at maximum speed *v_ab_* and the method is stable when Δ*x/*Δ*t* ≥ *v_ab_*. Unstable schemes arise when Δ*x/*Δ*t<v_ab_* because waves propagate more than one grid spacing in a period Δ*t*. However, for the 2D case one finds the stability criterion to be [86]

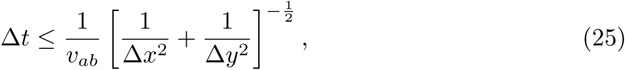

so, because Δ*x* =Δ*y*

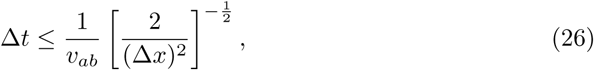

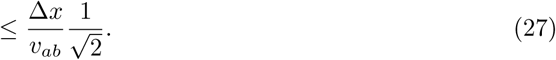

**Fig 5.**
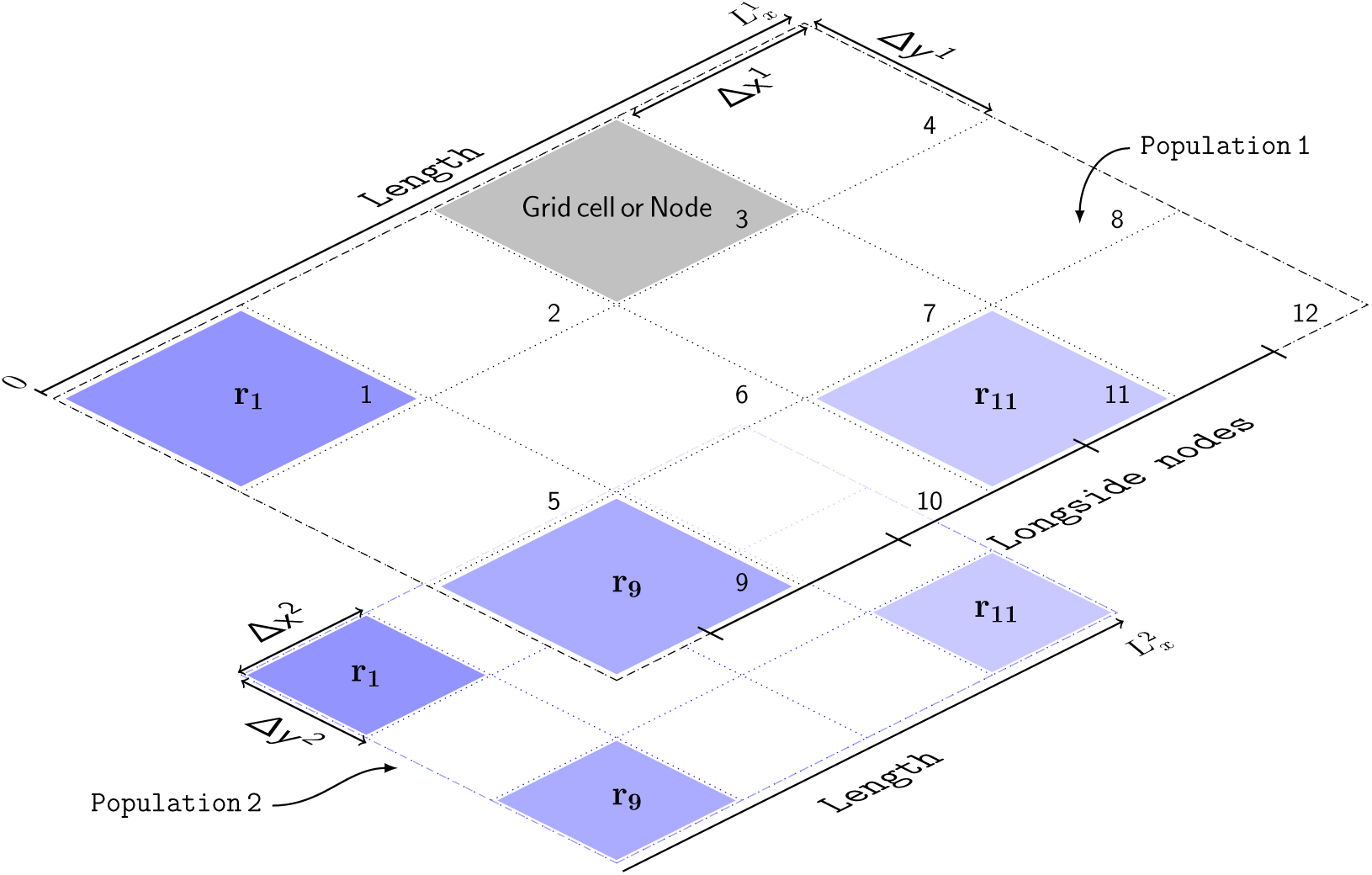
Schematic of the discretized spatial domain. The model has two populations: `Population 1` and `Population 2`. Geometrically, each population is represented by a grid of 12 nodes, which are labeled with integers. The grid is rectangular with dimensions 4 × 3 nodes. The number of nodes of the longest side is specified by `Longside nodes`. The physical size, *L_x_*, of each population is different. Thus, each node in `Population 1` has a linear size of Δ*x*^1^, and of Δ*x*^2^ in `Population 2`. Each spatial point (e.g., **r**_1_, **r**_9_, **r**_11_) is at the center of a grid cell. The subscript denotes the node index on this grid. Also, **r***_n_* denotes the actual position in the largest population; in the smallest population **r***_n_* denotes a rescaled physical dimension. Lastly, the borders of the grid are depicted with dashed lines to denote periodic boundary conditions (PBCs), which represent structures with planar geometry and toroidal topology.

Hence, considering all wave-type propagators, the maximum value of the Courant number *p*_max_ must satisfy

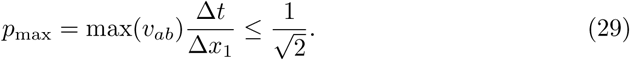

This condition is checked internally by *NFTsim* and if it is not satisfied, an error message is returned. Note that, in practice, one usually imposes a stricter condition to ensure the system has a margin of stability; e.g., in [2], the grid ratio was chosen so that *p*_max_ =0.1.

#### Explicit Difference Method and Boundary Conditions for the 2D Wave Equation

*NFTsim* uses an explicit central difference method [87] to solve Eq. (13), which represents axonal propagation of activity through the cortex or other structures with a significant spatial extent. Here, we present the explicit time stepping formula currently implemented to compute the next value of *ϕ_ab_* from past values of *ϕ_ab_* and *Q*_b_. The full derivation is in *Appendix S1*.

Equation (13) is the inhomogeneous damped wave equation, which can be simplified by making the substitutions

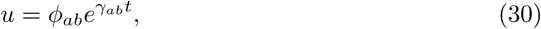

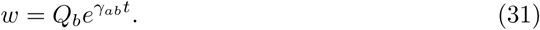

We then obtain the undamped wave equation

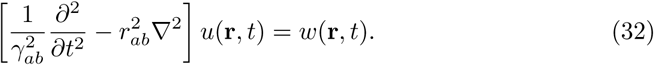

Note that this simplification only works for small values of Δ*t* because the exponential factors introduced in Eqs (30) and (31) diverge as Δ*t* →∞.

Then, the final time-stepping formula using an explicit central difference method for the 2D wave equation is

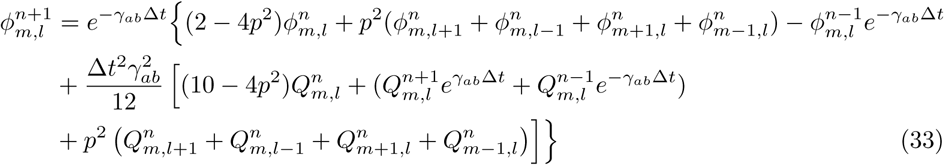
where the superscript *n* indexes time step; the first and second subscripts index space along the orthogonal *x* and *y* directions, respectively, except for the subscripts on *γ_ab_*; and *p* is the Courant number. In the Eq. (33), the positive exponential factors introduced in Eqs (30) and (31) are cancelled out and the algorithm actually evaluates negative exponential factors 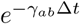 and 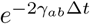.

Note that in Eq. (33), only five spatial points are required: the central point *m, l*; its horizontal neighbors *m* +1, *l* and *m* − 1, *l*; and, its vertical neighbors *m, l* + 1 and *m, l* − 1. This pattern is often referred to as a five-point stencil. There are alternative finite difference methods that use higher-order terms to approximate the derivatives and would require larger stencils (e.g., more neighboring points) [89]. It is usually better to increase the spatial resolution rather than the stencil complexity to obtain higher accuracy.

The finite difference scheme presented above is second-order accurate in space and time. This means that the rate at which the error between the discretized approximation and the exact continuum solution decreases to zero is 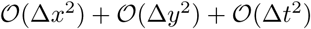. For instance, halving Δ*x*, Δ*y*, or Δ*t*, subject to Eq. (29) leads to a decrease of the error by a factor of four.

When solving partial differential equations on a finite spatial domain, one must specify boundary conditions for the simulations. *NFTsim* uses periodic boundary conditions (PBCs). This type of condition avoids boundary effects stemming from the finite size of a grid and avoids the perturbing influence of an artificial boundary like a reflective wall. In PBCs, opposite boundaries are treated as if they were physically connected, that is, the top of the grid is wrapped on to the bottom and the left of the grid on to the right.

The class `Stencil` has two main functions: (i) retrieving the five-point stencil pattern for every node in the grid; and, (ii) correctly copying the activity close to the boundaries of the domain at every time step to implement periodicity. To achieve this, `Stencil` operates on a grid of size (*N_x_* + 2) × (*N_y_* + 2). The additional ghost cells are used to store copies of the top and bottom rows and left and right columns of the grid.

Figure 6 illustrates a 4 × 4 grid with the additional ghost cells shaded in light blue labeled as `n, s, e, w` (i.e., north, south, east, west). The number in each grid cell represents its linear index – because the class `Stencil` accesses the elements of the two dimensional grid using a single subscript instead of two. The grid cells with prime, double-prime, and triple-prime indices are copies of the original cells with the same indices. These copies are used to implement PBCs along the vertical, horizontal, and diagonal directions, respectively. For instance, the cell 1′ is the vertical copy of cell 1; cell 1′ is the horizontal copy, and cell 1‴ is the diagonal copy. The diagonal copies are not used by the 5-point stencil, but would be used by a 9-point stencil [89].

**Fig 6.**
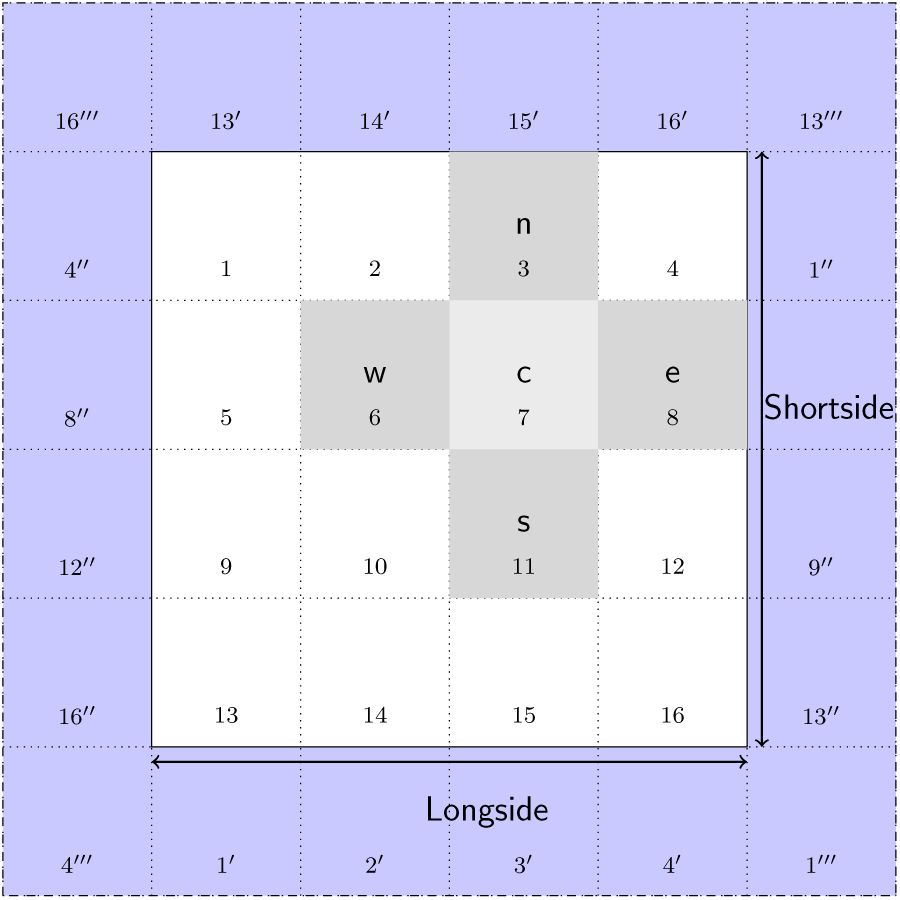
Schematic of the grid used by the class Stencil. This class retrieves the four nearest neighbors (labeled `n, s, e, w`) of a central point `c`. These five points define the pattern known as a five-point stencil. The cells in light blue are the ghost cells required to implement periodic boundary conditions. The prime, double-prime and triple prime indices represent copies of the corresponding indices in the vertical, horizontal and diagonal directions, respectively.

### Time Delays

For systems with time delays, *NFTsim* creates one history array of size *N* × *D_a_* per population, where *N* is the number of nodes as defined in previous sections. The delay depth *D_a_* of a history array is expressed as a number of integration steps.

To determine the exact value of *D_a_*, *NFTsim* checks every `Propagator` originating from *a*. Thus, the final delay depth for a given population *a* is

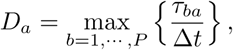
where *P* is the number of populations in the system; and, in the case that the parameter `Tau` represents spatially nonuniform time delays, *NFTsim* first selects the largest time delay over space

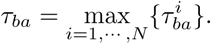

This produces history arrays with the minimum necessary delay depth for each population and thus is memory efficient.

Each `Propagator` stores an integer array with indices used to access the appropriate past activity of its origin population.

### Analysis and Visualization

*NFTsim* includes a Matlab module which provides ancillary tools to assist with running, analyzing and visualizing models. This package folder is called `+nf`. The available functions and a description of their functionality are summarized in Table 3.

The code snippet below uses some basic `nf` functions as an example of how users can interact with *NFTsim* directly from Matlab. The model is the same as the one specified in the configuration file `e-erps.conf` presented earlier, except that the timeseries of all the nodes in the grid are written to the output file. The simulation is executed via `nf.run()`. Once the output file is available `nf.read()` loads the simulated data into a Matlab structure.

Spatial patterns of activity and propagation of waves of activity across space can be visualized using the function `nf.movie()`

~~~
nf_ struct = nf.run(‘configs/e-erps-all-nodes.conf’)
nf.movie(nf _struct, ‘Propagator.1.phi’, 1)
~~~

Representative frames from the movie of waves propagating from stimulation sites are shown in Fig. 7(a) to Fig. 7(f). In each panel the mean spatial value of *ϕ_ee_*(*x, y, t*) at time *t* has been subtracted, so red and blue reflect positive and negative deviations, respectively, from the mean. The effects of PBCs can be appreciated from Fig. 7(c) onwards. In particular, in Fig. 7(c) the positive wave front propagating towards the left on the inset (*x* → 0) reappears at the right (*x* ≈ *L_x_*). In a similar way, Fig. 7(e), shows that the positive wavefront propagating along the *y*-axis results in a slight increase of the *ϕ_ee_* close to the boundaries.

The file used in this example is included in *NFTsim* and is also available in *Appendix S2*.

Furthermore, extracting and plotting the timeseries of a few nodes enable users to directly inspect the type of activity (e.g., healthy neural activity, evoked responses, or seizures). In this example, `nf.extract()` is used internally by `nf.plot timeseries()` to select the timeseries `Propagator.1.phi`.

~~~
these _nodes = {[1992:2008], [2089:2105]};
these _traces = {‘Propagator.1.phi’, ‘Propagator.1.phi’};
nf.plot_timeseries(nf _struct, these _traces, these _nodes, true)
~~~

Figure 8 shows the resulting plots generated with the code shown above. Each set of timeseries is centered around one of the stimulation sites. In Fig.8(a) the red curve is the axonal field at the site that received positive stimulation; and, in Fig.8(b), the blue line is the axonal field at the site that received negative inputs. The timeseries in gray above and below the colored curves are the axonal fields from neighboring sites along the *x*-direction. In these plots, the distance between the stimulation sites and neighboring sites increases vertically from the center to the top and bottom edges. The vertical dashed lines are not automatically produced by `nf.plot timseries`, but have been added to mark the onset time of the positive (red dashed) and negative (blue dashed) inputs, respectively.

**Fig 7.**
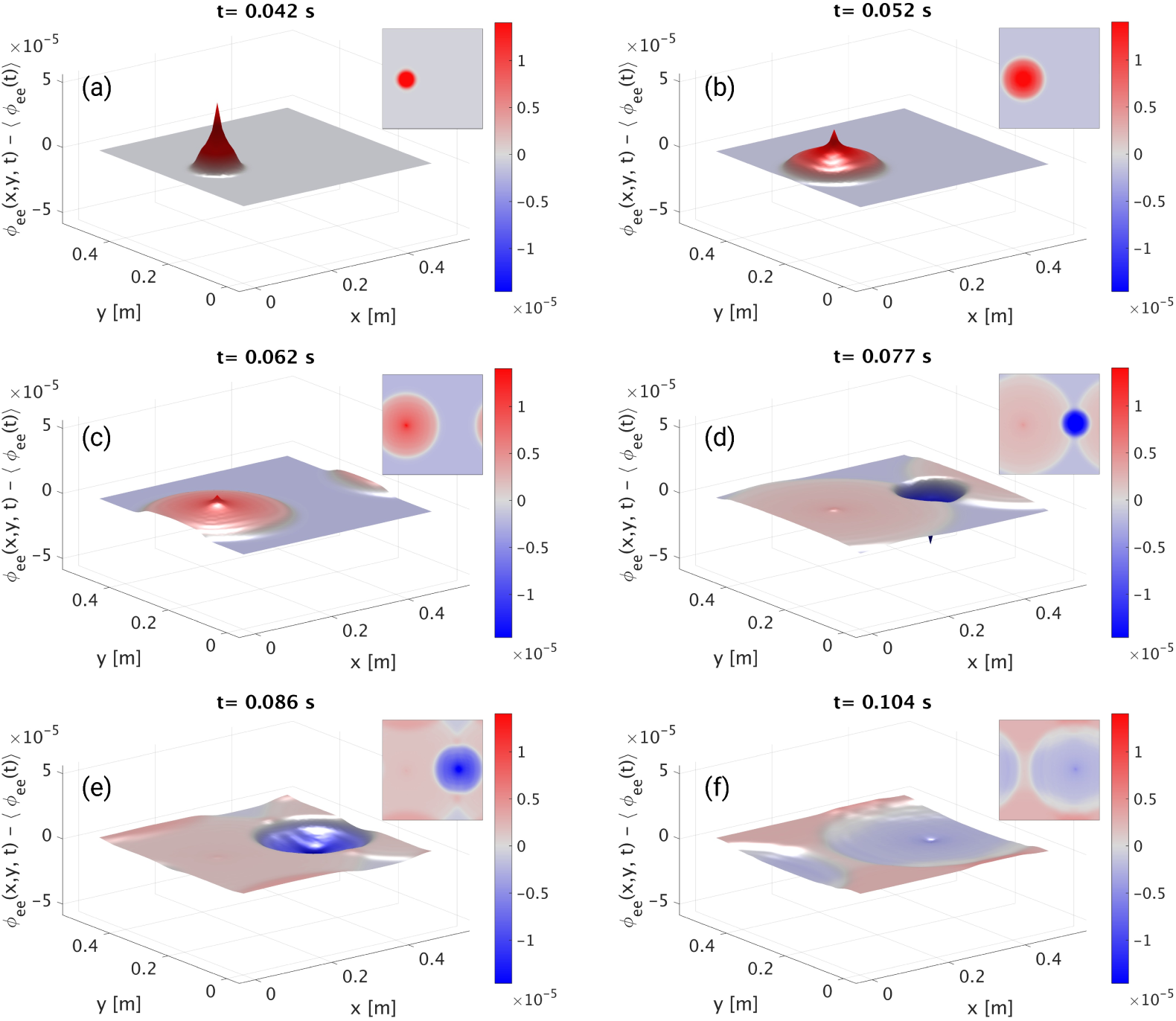
Neural activity of the model described in e-erps.conf. The cortical population is driven by two square pulses. The first pulse is positive, while the second pulse is negative. For illustrative purposes, in each panel the mean spatial value of *ϕ_ee_*(*x, y, t*) has been subtracted, so the color reflects deviations from the mean at that specific time. Each panel shows a surface plot of 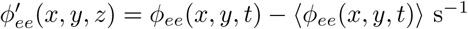 propagating radially outwards from the stimulation sites, and an inset with a planar view of the same quantity, at different times: **(a)** 42 ms; **(b)** 52 ms; **(c)** 62 ms; **(d)** 77 ms; **(e)** 86 ms; **(f)** 104 ms.

**Fig 8.**
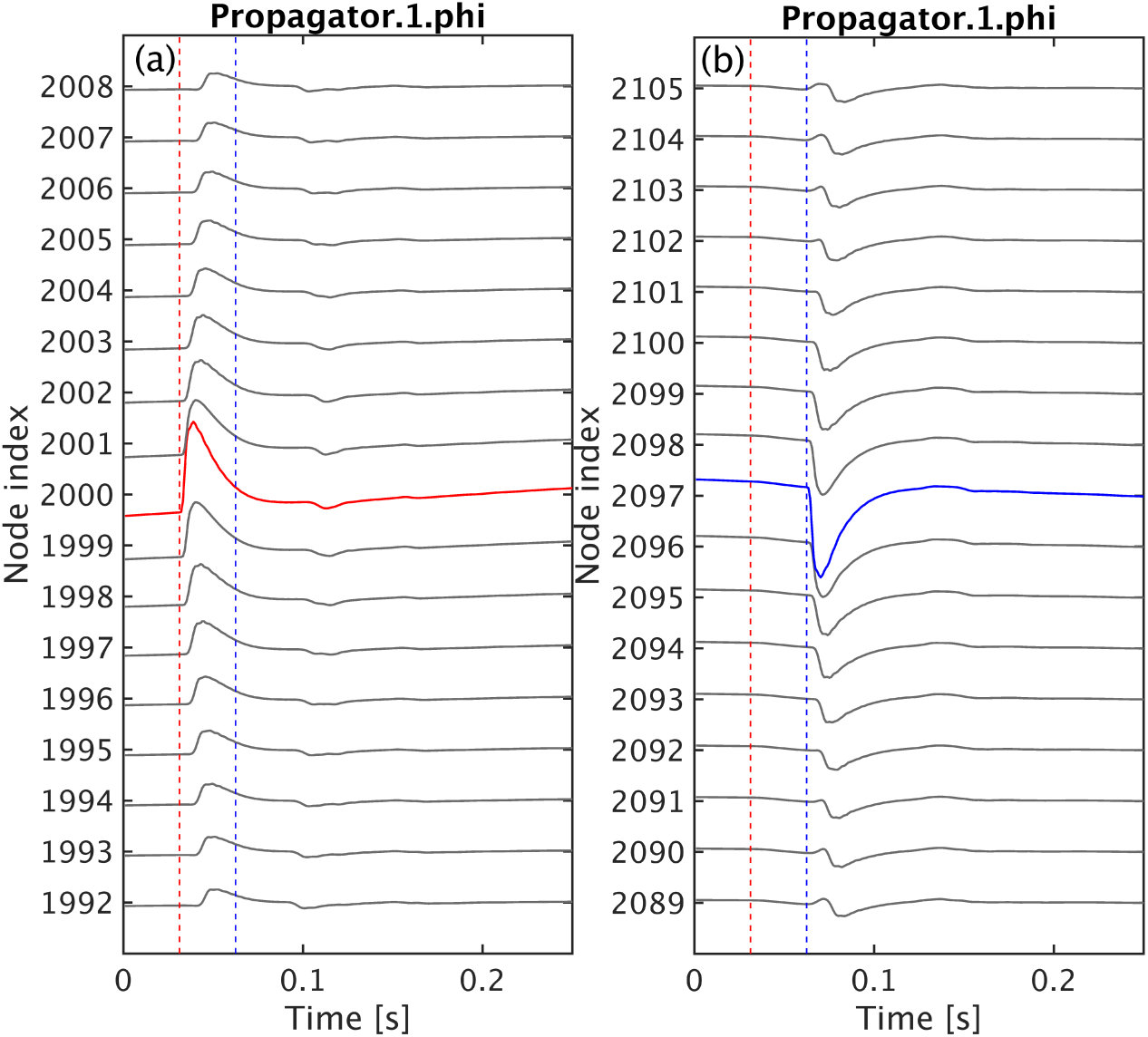
Timeseries of neural activity of the model described in e-erps.conf. The cortical population is driven by two temporal square pulses applied at the center of the grid as shown in Fig. 7. Here, we illustrate the timeseries of *ϕ_ee_* from a few nodes close to the vicinity of (gray lines) and at the stimulation sites. The vertical dashed lines mark the onset time of the positive (red dashed) and negative (blue dashed) stimulation inputs, respectively. **(a)** the axonal field at the site receiving the positive stimulus is highlighted in red while the time evolution of the same axonal field at neighbouring locations is shown as gray lines. **(b)** the axonal field at the site receiving the negative input is highlighted in blue.

Another important step is the calculation of the temporal power spectrum for a range of frequencies (in Hz), which is often compared to the power spectrum of experimental data. The power spectrum may also include multiple spatial modes for a range wave numbers (in m^−1^) and incorporate volume conduction or hemodynamic effects [100, 101] on measurement. A comparison between the linear analytic power spectrum and the numerical nonlinear power spectrum calculated with `nf.spatial spectrum()` is given as an example in *Standard Tests and Reproducibility*.

## Results and Applications

In this section we first present four exemplar systems that can be simulated using *NFTsim* and that have been previously studied in detail. Then, section *Observables and Diagnostics* briefly discusses the main observables that can be currently computed in *NFTsim* and how these have been used to predict a range of brain phenomena and compare to experimental results. In section *Standard Tests and Reproducibility*, we discuss how *NFTsim* could be used as a validation tool for neural field models and neural field simulators. Lastly, section *Benchmarks presents performance metrics and* practical information for users regarding average run times, memory usage, and storage required for typical simulations based on a neural field model of the corticothalamic system [29].

**Table 3.** Auxiliary functions available in the module `+nf`.

| Function | Description |
| --- | --- |
| extract | extracts time-series from an output structure |
| get_frequencies | returns the spatial frequencies |
| grid | reshapes output into a 3D array of shape $(L_{out}, N_x, N_y)$ |
| movie | generates a movie from a 3D array produced by `nf.grid` |
| plot_timeseries | plots vertically spaced timeseries of specific traces and nodes |
| read | reads an output file and returns a structure |
| report | prints information about a structure |
| run | runs a simulation from a configuration file |
| spatial_spectrum | computes spatiotemporal spectrum |
| spectrum | computes temporal spectrum |
| wavelet_spectrogram | computes wavelet spectrogram using a Morlet wavelet |

First column: names of the available Matlab functions. Second column: brief descriptions of what each function does. The variable $L_{out}$ is the number of time points in the output file.

### Exemplar Systems

The versatility of neural field theory and its concomitant implementation in *NFTsim* allow for the investigation of an unlimited variety of specific models and parameter sets. In this section we present a few illustrative cases, which have been thoroughly described elsewhere, along with some of their applications [1, 2, 6, 27–31, 33–39, 71]. Their respective configuration files are included in *NFTsim*.

The most general corticothalamic model considered here includes populations with long-, mid-, and short-range connections in the cortex, the specific and reticular nuclei in the thalamus, and external inputs. We indicate how components of this model can be successively deleted to obtain a family of models suited to simpler applications in corticothalamic and cortical systems. In what follows we label specific models according to the internal populations they include. The first system, called EMIRS, includes five different populations of neurons: cortical excitatory pyramidal (*e*), excitatory mid-range (*m*) and inhibitory (*i*) populations; internal thalamic reticular nucleus (*r*), relay specific nucleus (*s*), whereas the simplest case is of a system with a single excitatory population (*e*). There is also always at least one external population that provides inputs (often labeled either *x* or *n*). *NFTsim* provides a number of external input types such as sinusoids (in space and time), pulses, and Gaussian white noise. For instance, these inputs could be from excitatory neurons of the auditory pathway, which transmit signals from the cochlea to the thalamus [73]; or, they could be artificial external stimulation like Transcranial Magnetic Stimulation [38].

Figure 9 shows schematics of the illustrative neural field models described here. The EMIRS corticothalamic model displayed in Fig. 9(a) includes three cortical populations (*e*, *m*, and *i*) and two thalamic populations (*r* and *s*), with intracortical, intrathalamic, and corticothalamic connections.

The EIRS corticothalamic model is obtained by deleting the population *m* from Fig. 9(a) to obtain Fig. 9(b). Physically, this deletion corresponds to describing the effects of the mid-range population, whose axonal range is of the order of a few millimeters, as part of the short-range *i* population [1]. In this case, the excitatory effect partly cancels inhibition to give a weaker, net effect from this compound population, which includes the effects of both short-range excitatory and inhibitory interneurons. This model has been successfully applied to investigate a wide range of phenomena [31, 43, 72] (see *Introduction*). The model has five distinct populations of neurons: four internal and one external.

In the purely cortical EI model of Fig. 9(c), thalamic dynamics are deleted and *ϕ_es_* = *ϕ_is_* is assumed to replace *ϕ_sn_* as the external input to an isolated cortex. The basic EI model includes external inputs to two cortical populations (*e* and *i*), and both intracortical and corticocortical feedback are represented. This model is a starting point for understanding more elaborate neural fields models of the cortex (e.g., modeling distinct layers within the gray matter [35, 73]). Delays in the propagation of signals within neurons are due to synaptodendritic, soma, and axonal dynamics. However, in this model there are no long-range delays like those from the thalamus to the cortex. An extensive description and analysis of this model are given elsewhere [2, 32, 74], including emergence of gamma rhythm [72] and integration of cholinergic modulation [75]. Finally, the excitatory-only E model in Fig. 9(d) omits cortical inhibitory effects. This neural field model is the simplest system we consider that can be simulated in *NFTsim* and has been used as the central example throughout this work.

### Observables and Diagnostics

Brain phenomena including the alpha rhythm [31, 34], age-related changes to the physiology of the brain [27], evoked response potentials [28, 35, 73], and seizure dynamics [1, 5, 36, 71], can be measured noninvasively via EEG. In these studies, the fields of activity of the excitatory cortical population *ϕ_ee_* have been used to approximate EEG signals measured from the surface of the scalp [50, 90] and constitute one of the main biophysical observables comparable to experimental EEG data.

For the reasons mentioned above, the neural activity produced by *NFTsim* closely resembles the electrical activity measured by EEG and ECoG up to a dimensional constant (i.e., translate units of rate (s^−1^) into voltages).

Another tool traditionally used to detect various waking and sleep stages [6, 29] is the EEG power spectrum [50]. In calculating scalp EEG spectra (rather than intracranial ones), one must consider filtering due to volume conduction by the cerebrospinal fluid, skull, and scalp [50, 90]. The calculation of the power spectrum including volume conduction filtering is implemented as a spatial filter in `nf.spatial spectrum`. The smoothed EEG timeseries can be obtained by inverse Fourier transforming the filtered power spectrum. In the case of ECoG, the spatial filtering due to volume conduction should not be applied.

It is important to notice, though, that the neural activity of different cortical and subcortical populations can be used to predict other relevant electrical signals such as local field potentials (LFPs), and stereoencephalography (SEEG); magnetic signals such as MEG; metabolic-related signals like fMRI [91] or indirect fluorescence signals like those recorded via voltage sensitive dyes imaging (VSDI) [92]. Note that conversion of *NFTsim* outputs to the desired neuroimaging modality signals still requires additional modeling steps, including a description of the causal relationship and physiological couplings between the sources (i.e., the spatiotemporal fields of neural activity stemming from multiple populations) and the effective biophysical quantity measured in experiments [93, 95, 96, 99–101].

**Fig 9.**
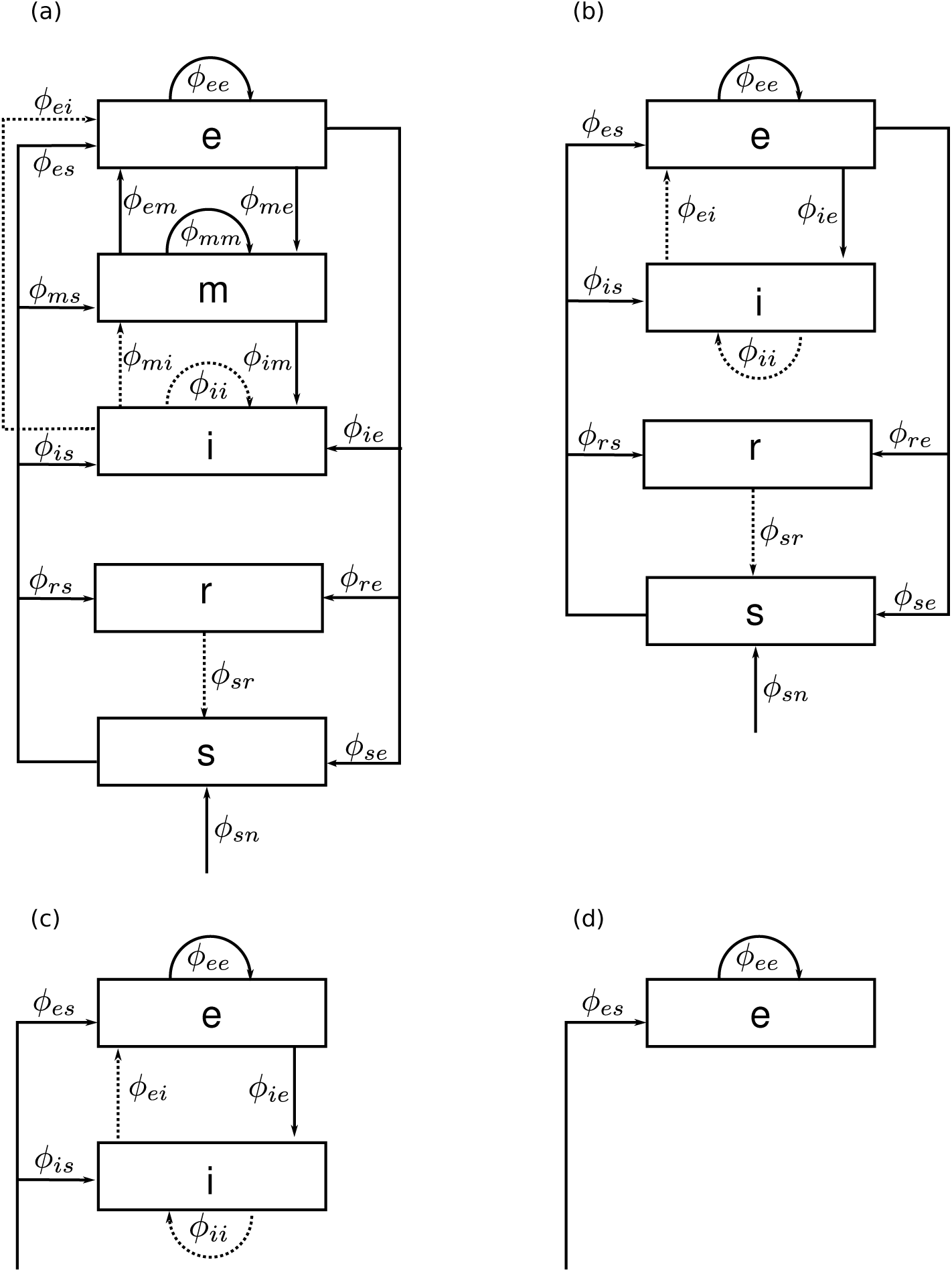
Schematic of four representative neural field models. The quantities *ϕ_ab_* are the fields propagating to population *a* from population *b*. Dashed lines represent inhibitory connections. **(a)** Corticothalamic model including excitatory (*e*), mid-range (*m*), inhibitory (*i*), reticular (*r*), specific relay (*s*) and external non-specific (*n*) populations. **(b)** Corticothalamic model including excitatory (*e*), inhibitory (*i*), reticular (*r*), specific relay (*s*), and external (*n*) populations. **(c)** Cortical model comprising excitatory (*e*) and inhibitory (*i*) cortical populations plus an external input field from a subcortical population (*s*). **(d)** Purely excitatory (*e*) cortical model with input from a subcortical population (*s*).

### Standard Tests and Reproducibility

Standard tests are a set of benchmarks used evaluate and compare disparate numerical implementations of similar neurobiological models [97]. There are very few such tests in computational neuroscience [98] and the ones currently available are only for single-cell models. To the best of the authors’ knowledge, there are no published standard tests for mesoscopic models such as neural fields. Thus, there is a huge void regarding quality assessment of scientific software for continuum models of brain dynamics.

*NFTsim* provides a reference framework for standard tests for implementations of neural field models because its methods have been verified with analytic results; and, the linear analytic closed form solutions upon which the code is based have been extensively validated with experiments, as discussed in the Introduction. For example, in Fig. 10 we reproduce the results presented in Fig. 2 from [6]. This plot shows a comparison between the linear analytic power spectrum (dashed line) and the spectrum computed from *NFTsim* simulations. Both spectra agree within the range of 0.1–45 Hz with a root-mean-square error of approximately 6 × 10^−20^. *NFTsim*’s default `eirs-corticothalamic.conf` is used with the parameters from [6], which we do not repeat here because the original configuration files are also included as part of the library package. The power spectrum is calculated using the `nf.spatial spectrum()` function.

Furthermore, the *NFTsim*’s methods and implementation have also been directly validated by experimental data for nonlinear dynamics, notably neural activity corresponding to seizures [36] and sleep spindles [6].

**Fig 10.**
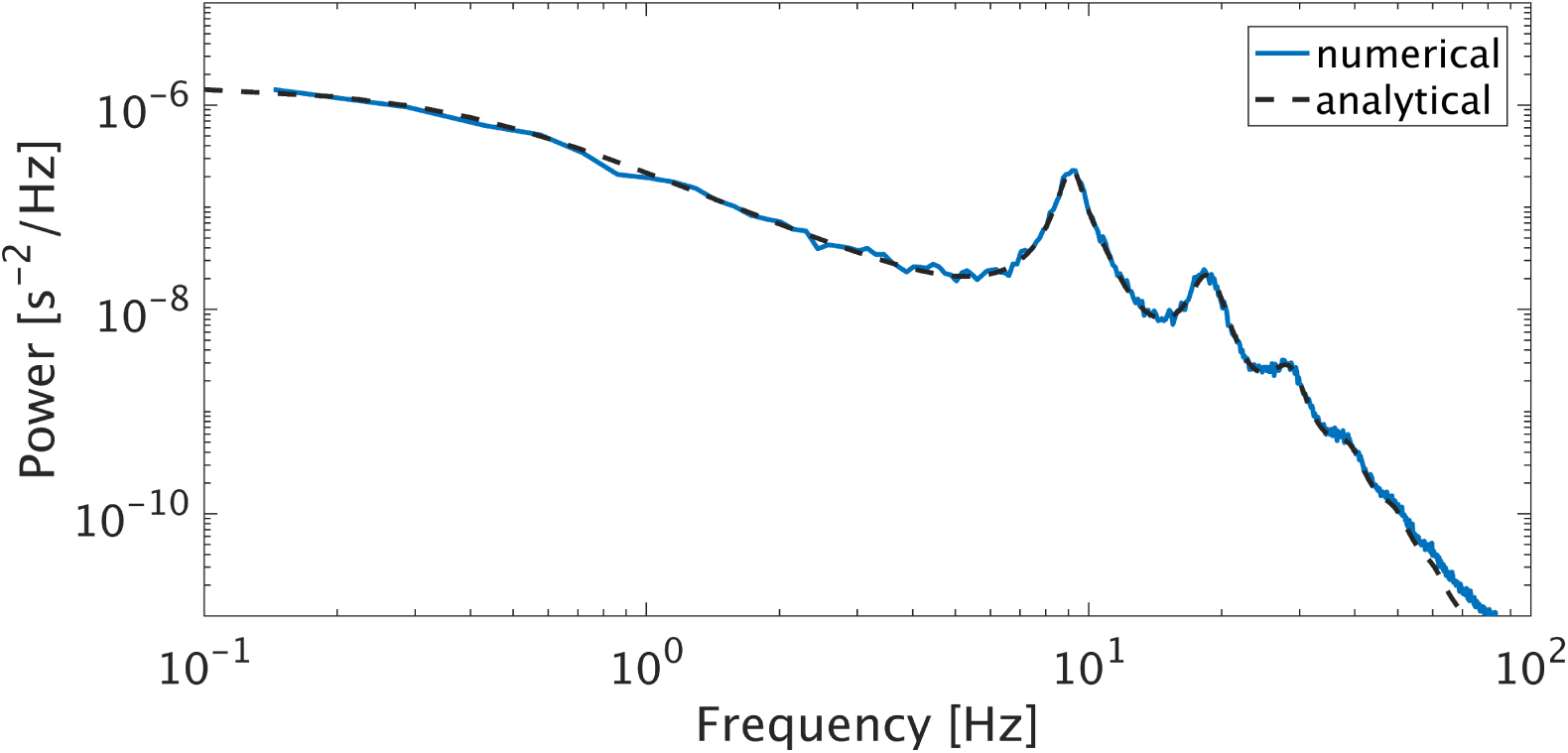
Comparison of analytic and numeric EEG power spectra in the corticothalamic system. The dynamics of the EIRS model were simulated using the wake parameters from [6] for their Figure 2. The linear analytic spectrum (black dashed solid) is compared against the spectra computed from simulations (solid line).

### Benchmarks

*NFTsim* provides a tool for semi-automated benchmarking. Timing and configuration information for simulation runs are stored in a comma-separated value (csv) file that can be processed at a later stage.

Invoking the script

~~~
nf _benchmarks
~~~

without any arguments will run all the configuration files in the `benchmarks/` directory once. We provide ten default configuration files that run in a total of under 700 s on a desktop computer. Example results for specific hardware are given below. These files are based on the corticothalamic model and are representative of typical simulation scenarios. With nf_benchmarks users can also:

i. benchmark a specific configuration file ~~~
nf_benchmarks <config_filename>
~~~
ii. benchmark a specific configuration file multiple times (e.g., 8 times in this example) ~~~
nf_benchmarks --num - trials 8 <config_filename>
~~~
iii. benchmark a specific configuration file with output written to memory instead of disk (this only works under Linux) ~~~
1 nf _benchmarks --to-mem <config _filename>
~~~
iv. benchmark a specific configuration file using a non-default compiler ~~~
1 nf _benchmarks --clang <config_ filename>
~~~

In *NFTsim* propagating fields are followed via partial differential equations, so the main contributions to the runtime *T* are (i) the number of grid cells *N*; (ii) advancing a maximum of *P* ^2^ fields, between *P* populations, on the *N* cells; (iii) the length of the simulation in integration steps *L* = *T*_sim_/Δ*t*; and, (iv) the size of the output *O* written to file. So,

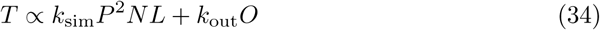
where the coefficients *k*_sim_ and *k*_out_ depend on the hardware architecture. The output size *O* depends on the product of the total number of variables (*W*), the number of grid cells (*N*_out_) and the total output time points [*L*_out_ =(*T*_sim_ − *T*_start_)/Δ*t*_out_] requested in the configuration file.

For large *O*, the runtime is dominated by writing operations. This overhead is expected for two reasons: (i) a simulated data sample is written to disk every Δ*t*_out_, which takes additional time; and, (ii) writing the output to a text file requires conversion of numbers to text. Despite the runtime overhead this last point entails, text files are a convenient format to store the output because they are easier to debug than binary files.

The required memory, *M*, used by a *NFTsim* process is dominated by the number of grid points *N* and the history arrays of *P* internal populations with delay depth *D_a_*, which is the number of integration steps for a signal to propagate to the target population from the source population. So,

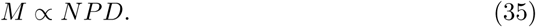

Table 4 summarizes the simulation parameters that determine runtime and memory usage of a *NFTsim* process, including those which are not directly specified in a configuration file.

To assess *NFTsim*’s performance, we select the corticothalamic model presented in earlier sections, with the parameters taken from previously published work [6] and thus considered a typical simulation use case.

The simulation length and integration time step are held constant at 16 s and Δ*t* =2^−14^ s ≈ 10^−4^ s, respectively. So, the only varying parameter that affects the runtime and storage is `Nodes` (*N*). The choice of this integration time step size is such that is sufficiently small to resolve high frequency oscillations and to satisfy the Courant condition for numerical stability for a range of discretization values between 3 mm < Δx< 50 mm. The Courant number ranges between 0.014 *< p* < 0.15 for a fixed velocity *v_ab_* =10 m s^−1^.

**Table 4.** NFTsim simulation and output size parameters, and runtime and memory usage symbols.

| Symbol | Description | Parameter in configuration file | Units |
| --- | --- | --- | --- |
| $T_{sim}$ | Simulation length | Time | s |
| $\Delta t$ | Integration time step | Deltat | s |
| $L$ | Number of time steps | - | - |
| $P$ | Number of populations | - | - |
| $D$ | Delay depth | - | - |
| $O$ | Output size | - | - |
| $N_{out}$ | Number of output nodes | Node | - |
| $\Delta t_{out}$ | Output sampling period | Interval | s |
| $T_{start}$ | Output start time | Start | s |
| $W$ | Number of output variables | - | - |
| $L_{out}$ | Number of output time points | - | - |
| $M$ | Memory used | - | byte |
| $T$ | Runtime | - | s |

First column: symbols used in this work to identify either the parameters specified in a configuration file, or variables associated with runtime and memory usage. Second column: description of the variable or parameter. Third column: parameter names used in configuration files. The symbol – means the parameter is not specified directly in a configuration file. Fourth column: SI units. Here, the symbol – means the quantity is dimensionless.

Two groups of simulations were run. The first group, *G_no_*, runs the simulation and only writes a copy of the configuration file to the output file. The subscript *no* means `no output`. In this case the runtime represents the effective time spent executing a simulation without the time overhead due to writing operations. From Eq. (34), the group *G_no_* has *k*_out_ ≈ 0. The second group of simulations *G_wo_* consists of identical simulations to those of *G_no_*, except that all the model variables (firing rate, voltages, fields, coupling strengths), for all the nodes, sampled at 512 Hz, are written to a file in the hard disk.

Approximate runtimes and memory usage are measured using tools available on Linux systems. The computer used for the benchmarks has Red Hat Enterprise Linux (RHEL) 6.9 as operating system, GNU Compiler collection (gcc) 4.9.2 as the default compiler, a 3.50 GHz Intel i5–4690 processor and 8GB of RAM.

Table 5 presents the benchmark results for different grid sizes and shows that the runtimes scale linearly as a function of the number of nodes with *k*_sim_ ≈ 0.15 s for the simulation group *G_no_* and and *k*_sim_ ≈ *k*_out_ ≈ 0.15 s for group *G_wo_*. From these results, we conclude that in order to produce one minute worth of data sampled at a rate typically used in clinical EEG recordings, *NFTsim* takes about four minutes to run the simulation and write the output to disk. Thus, *NFTsim*’s simulation length to real-time data length ratio (*T*_sim_*/T*_real_) for EEG-compatible outputs is approximately 4. To reduce this ratio users can decrease the size of the output *O*, by writing only a few relevant variables to disk.

While these benchmarks offer a narrow view of *NFTsim*’s performance, they are a valuable practical tool for users and provide: (i) estimates of resources required to run simulations; and, (ii) a guide to make informed decisions between the execution runtimes and accuracy (i.e., decreasing the spatial resolution and/or the time step).

**Table 5.** Benchmarks for different grid sizes using NFTsim v.0.1.5.

| Nodes | Storage [GB] | Memory [MB] | Runtime $G_{no}$ [s] | Runtime $G_{wo}$ [s] |
| --- | --- | --- | --- | --- |
| 144 | 1.1 | 5 | 24 | 44 |
| 256 | 1.9 | 6 | 40 | 79 |
| 1024 | 7.3 | 16 | 161 | 302 |
| 4096 | 29 | 51 | 646 | 1208 |
| 16384 | 116 | 200 | 2601 | 4908 |

The value of *Nodes* is reported on the first column. The second column informs the storage size of the output file with all the model variables for each node (firing rate, voltages, fields, coupling strengths) sampled at $1/\Delta t_{out} = 512$ Hz for the group of simulations $G_{wo}$ . The third column presents the total memory usage of the process. Reported runtimes on the fourth column and fifth columns are the time elapsed in seconds. The values on the fourth column correspond to simulations with no output (no) written to the output file. The fifth column corresponds to the running times of simulations for which all the model variables for every node of the grid are written to the output file (wo). The same corticothalamic model is used in every simulation with $T_{sim} = 16$ s, and $\Delta t = 2^{-14}$ s.

## Conclusions, Availability, and Future Directions

We have introduced *NFTsim*, a user-ready, extensible and portable suite for numerical simulations of neural activity based on neural field models expressed in differential form. *NFTsim* is based on the well established framework of neural field theory [2] and has been validated with both analytic solutions and experimental data. Thus, when working with new models and simulations users can use analytic solutions as a way to validate their results.

Written in C++, *NFTsim* has been tested on a range of Linux distributions (RHEL 6.9, RHEL 7.4, OpenSUSE 13.2, OpenSUSE 42.2). Current users have reported compatibility with OS X 10.11 and mac OS 10.12 in conjunction with the CLANG compiler, provided that the C++11 standard is supported. *NFTsim* has not been tested under Microsoft Windows.

The output of *NFTsim* is written to a plain text file and ancillary modules written in Matlab contain functions to assist in simulation execution, quick analysis and visualization of the results. *NFTsim* thus provides an efficient solution to simulating various continuum spatiotemporal models including spatially uniform (homogeneous) and nonuniform (inhomogeneous) neural field models [80]; systems with heterogeneous time delays between populations [34]; and, the selected format for data storage is simple enough that enables users to choose from a broad selection of tools to perform further analysis and visualization. The development of *NFTsim* follows essential practices of modern open-source scientific software development [98] such as:

i. The code is licensed under the Apache 2.0 license.
ii. Our code sources are hosted on Github: https://github.com/BrainDynamicsUSYD/nftsim.
iii. We use pull requests to review new features and bug fixes.
iv. Our users can open issues reporting bugs and/or other problems they encounter.
v. The developer documentation is produced using Doxygen [103].
vi. A separate manual is provided for end-users.
vii. Releases are tagged, so users can refer to and download continuously improved versions of the code that are considered stable and tested. For instance, for this paper, we have used v.0.1.5.

Most notably, the activity from neural populations can be used to calculate biophysical signals such as LFP, ECoG, or EEG signals, the latter being the most commonly found in previous studies. Other forms of biophysical observables, such as fMRI and VSDI may also be implemented, but require additional modeling work to define how the electrical activity relates to the corresponding measurements (e.g., oxygen consumption, blood flow changes or fluorescence). Further physical effects can be implemented as a part of postprocessing modules like `+nf`.

Due to its flexibility and generality, *NFTsim* allows for a systematic study of both healthy and unhealthy brain function. For instance, in [6] the authors used simulations of a full nonlinear EIRS model for parameter values representing typical sleep spindle oscillations. They found that the numerical nonlinear power spectrum had an additional harmonic peak that was neither present in the linear EIRS model nor it was predicted by the analytic linearized power spectrum. This study clearly demonstrated that *NFTsim*’s flexibility allowed for the investigation of nonlinearities, introducing them one at the time in different neural populations. This enabled the authors to determine which anatomical structures and physiological mechanisms were responsible for the dynamics observed in experiments.

Due to its modularity, *NFTsim* is extensible and can accommodate new features presented in theoretical work on neural fields. In fact, a tool like *NFTsim* is essential for the study of nonlinearities and connectivities configurations that do not necessarily follow the random connectivity approximation [2, 51, 108] or are not spatially homogeneous or constant over time. For instance, [71] explored the mechanisms of seizures by incorporating slow currents modulating the bursting behavior of the reticular nucleus in the corticothalamic (EIRS) model; while [38] incorporated a model of synaptic plasticity to the purely excitatory subsystem. These two mechanisms are already implemented in the current version of *NFTsim*. However, further investigation and development work is required before implementing a general mechanism of parameter modulation, which would allow for the study different types and functional forms of neural feedbacks [61, 102].

We remind potential users that *NFTsim*, as any scientific software, should not be used blindly. As a minimal requirement, users should check that:

i. The integration time step is small enough to resolve the simulated dynamics correctly, especially if the system exhibits chaotic and bursting dynamics;
ii. The parameter `Interval`, which effectively subsamples the timeseries written to disk, is an integer multiple of the time step; and, is small enough so as to avoid temporal aliasing if there are signals with high-frequency content (e.g., the effective sampling frequency of the signal written to disk (1/`Interval`) is sufficient to respect Nyquist’s sampling theorem.
iii. The integration time step is small enough to respect the Courant conditions. If this condition is not met the code throws an error. A way to select an appropriate value of `Deltat` would by running the simulation with increased or decreased time steps to check for stability and convergence of the solutions to a limiting case.
iv. Setting parameters such as `Deltat`, `Interval` and `Nodes` as multiples or submultiples of powers of two (*NFTsim*’s default values), minimizes numerical errors due to the inherent limitations of representing floating point numbers on a computer. In addition, other advantages of using powers of two are (i) achieving optimal performance of FFT algorithms when applied to the output timeseries (1 second of data will have a power-of-two number of samples); and, (ii) avoiding zero-padding which is a frequent default behavior of FFT algorithms. However, users are not obliged to use *NFTsim*’s default values. They can select any value and simulations will be executed.
v. Artifacts of periodicity introduced by PBCs as illustrated in Fig. 7 are avoided. This can be achieved by setting the grid’s area larger than that of the actual physical system under consideration. In this scenario, waves propagating from the region of interest towards the right edge of the grid would die off before being reintroduced on the left edge. This approximation would be close to the solution in the absence of artificial boundaries in which the region of interest has infinite size; or to absorbing boundary conditions (ABCs).

Note that PBCs may be preferable over absorbing BCs in scenarios when one is interested in studying wave-wave interactions. However, in neural field models that approximate a small patch of cortex such as the primary visual cortex V1, in which waves of activity propagate away from a source point of stimulation [96], then absorbing BCs would likely be more appropriate.

In the present work we have concentrated mainly on a high-level description of the software and presented examples for which model parameters are assumed to be spatially uniform. *NFTsim* already accepts spatial variations in many parameters, although more development work needs to be done to provide general mechanisms of parameter variation.

Note that *NFTsim* solves differential equations. It does not solve integrodifferential equations like those of Nichols and Hutt’s Neural Field Simulator [8]. It is important to notice that not all neural field expressed in integrodifferential form can be expressed in differential form. On the other hand, neural fields expressed in differential form can be expressed in integrodiffrential form and can be solved in *NFTsim*.

As mentioned in *Classes and their Interactions*, *NFTsim* currently has Gaussian white noise in its collection of external driving signals because in the literature [1, 2,4, 8,28–31, 39, 65, 104–107] neural field models are typically either initialized or driven by a signal with a flat (white) power spectrum. These inputs correspond to the activity of other brain structures that are not explicitly modeled in the equations. In the case of the corticothalamic system, these inputs may represent background activity from the brain stem. In the case of a purely cortical model, these inputs could represent the combined activity from the thalamus and other structures. It is important to notice that external inputs with a broad white spectrum enter the NFT differential equations in exactly the same way as more coherent stimulus such as a sine wave would, as such, standard numerical methods (RK4) are employed.

However, there are several limitations that make this type of signal a poor choice. The first limitation is that idealized continuous noise is not physically realistic because it has an infinite bandwidth and infinite power. The second limitation is that in computer simulations, where continuous models are inevitably discretized, the bandwidth of a white noise signal depends on the size of the discretization. This dependence implies that if either the time step or the spatial step are reduced, the bandwidth increases and as a result a white noise signal has additional modes (i.e., frequency components). One can use a scaling parameter to adjust the overall power of the discretized driving signal [6, 87]. This scaling has no effect on the resulting spectral shape that is often compared to EEG [6]. For these reasons, it is necessary to incorporate a new type of stimuli that has a white spectrum but that is differentiable in time and space; and its spectral profile does not change under changes of the discretization.

Future work will extend *NFTsim* scientific features by including (i) a new bandlimited noise generation to render the inputs even more biologically realistic; (ii) generalized mechanisms of spatiotemporal variations for different model parameters and variables; (iii) generalized mechanisms of neuromodulation; (iv) absorbing boundary conditions; and, (v) spherical topology. In addition, a number of technical enhancements will be made such as (i) implement support for output binary files; and (ii) extend and automate unit test coverage to ensure that new additions to the code do not break previous functionality.

## Supporting information

### Appendix S1. Discretization of the wave equation

In this section we describe the discretization of the wave equation. This method allows us to obtain an equation to advance each field *ϕ_ab_* from *t* to *t* +Δ*t*. We remind the reader that the equation relating the field *ϕ_ab_*(**r**, *t*) to the driving signal *Q_b_*(**r**, *t*) is

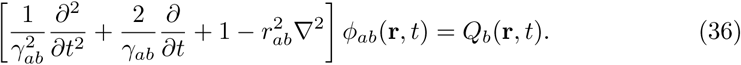

This equation is a damped wave equation for *ϕ_ab_*(**r**, *t*) with source *Q_b_*(**r**, *t*). The damping is introduced via the first-order derivative term in the same way friction forces enter a vibrating mechanical system; and, by the third term in Eq. (36). This equation can be simplified by making the following substitutions

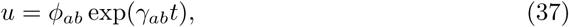
and

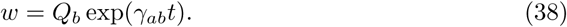

We then obtain the undamped wave equation

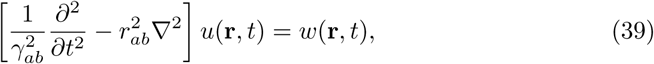

To solve this differential equation numerically, we replace the temporal and spatial derivatives with finite central difference approximations on a discretized domain. The derivation presented in the following paragraphs solves the Eq. (39) by using explicit methods, that is, the next value of *ϕ_ab_* is computed from known past values of *u* and *w* and all future time terms appear on the same side of the time stepping equation.

Consider first the term *∂*^2^*/∂t*^2^ in Eq. (39), and let the superscripts *n* index time in units of *k* =Δ*t*. We can use a Taylor expansion to write

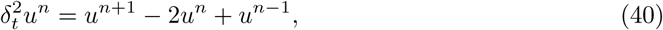

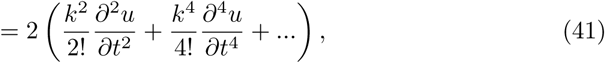

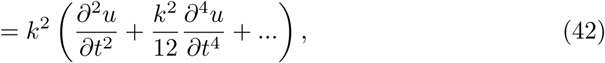
where 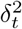 is the second order central difference operator in time; and, *u^n^*^+1^ is the future term we are interested in calculating. Combining Eqs. (40) and (42) yields

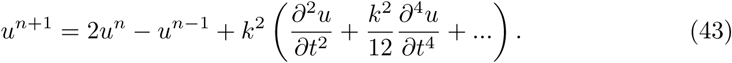

Note that this approximation is 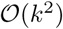 accurate in time because we use a second order central difference formula to approximate the second order derivative. So, the error is proportional to the square of *k*. In a similar way, the second order centered finite difference approximation for the second order spatial derivatives are

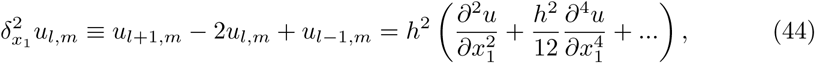

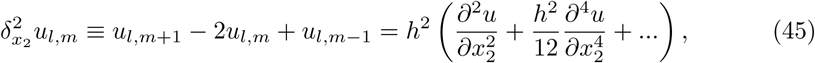
where h = ∆*x*_1_= ∆*x*_2_ is the grid spacing and the subscripts m and l index grid points in the orthogonal *x*_1_ and *x*_2_ directions, respectively. The error of the centered difference scheme used here is 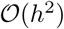. We also use:

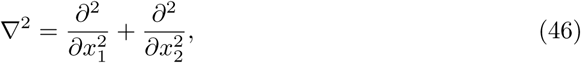

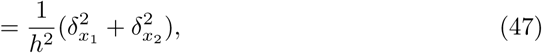
and

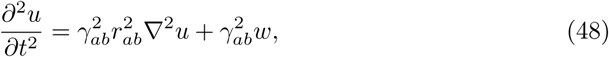

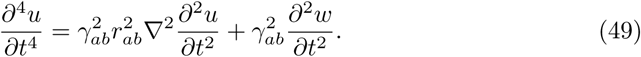

We then substitute Eqs (48) and (49) into Eq. (43) and obtain

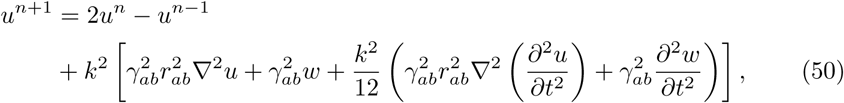
and further substitute the term 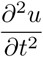 in Eq. (50) for the right hand side of Eq. (48)

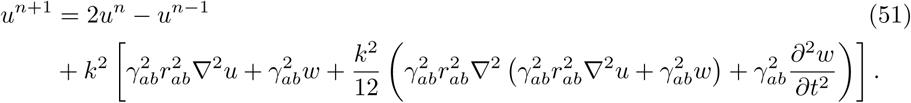

By rearranging the terms in Eq. (51) we can express *u^n^*^+1^ in terms of *u* and *w*

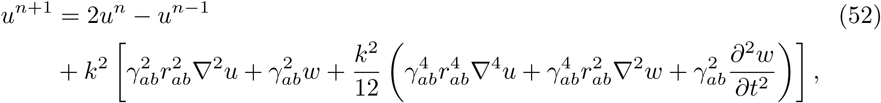

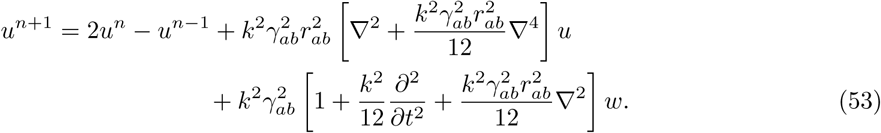

We now omit the terms involving ∇^4^ since a second order approximation is enough, giving

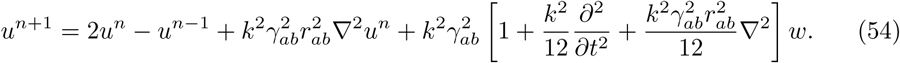

Next we replace ∇^2^ by the approximations defined in Eq. (47) to obtain

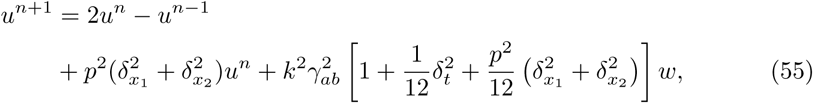
where *p* ≡ *p_ab_* = *kγ_ab_r_ab_/h* is the Courant number and is equivalent to Eq. (29). Next, we replace the second order difference operators 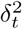, 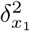, and 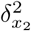 to obtain an explicit solution to compute the next value in time of *u_m,l_*:

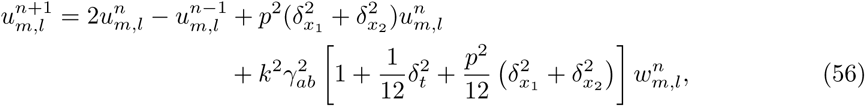

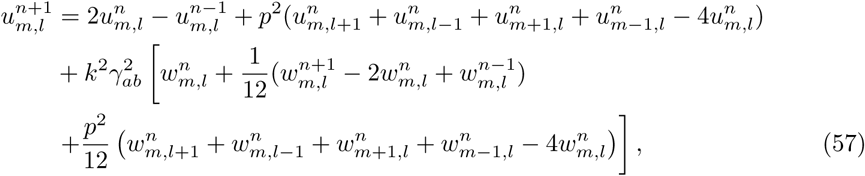

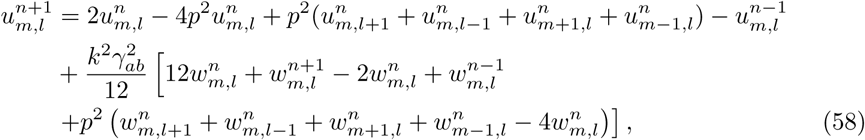

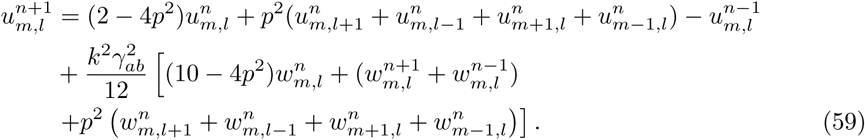

From Eqs (37) and (38), 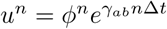 and 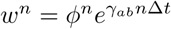. Also, for a single simulation step, the current state is centered at *t* = 0 and thus indexed by *n* = 0; the next and previous states are ±1 step away, or equivalently ±Δ*t*. Then, *n* + 1 denotes time Δ*t* and *n* − 1 denotes time −Δ*t*. Therefore we define the following substitutions

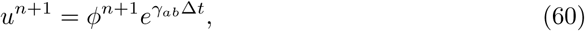

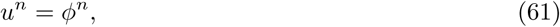

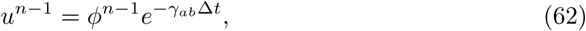

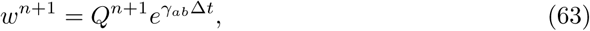

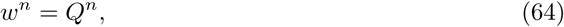

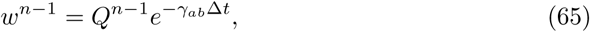

The spatial indices are omitted for compactness but can take the values {*m, m* ± 1} and {*l, l* ± 1}. Hence, Eq. (59) can be expressed in terms of *ϕ* and *Q* as

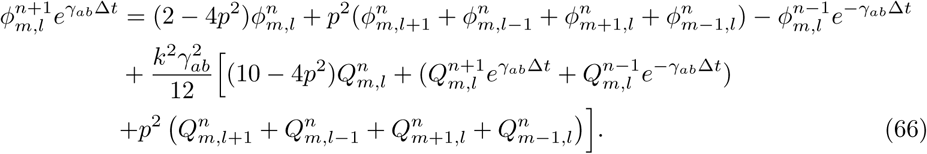

Finally, upon multiplying both sides of Eq. (66) by 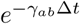 one finds

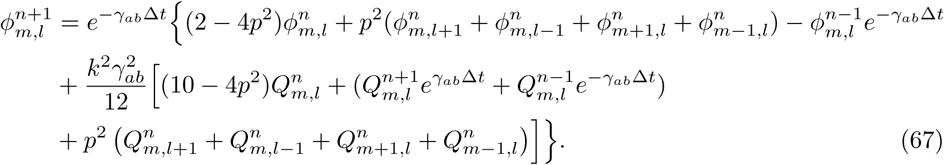

Eq. (67) is the formula to advance an axonal field *ϕ_ab_* one time step based on its current state (*n*) and previous state (*n* − 1) when *ϕ_ab_* is governed by Eq. (13).

### Appendix S2. Configuration file used in Analysis and Visualization

~~~
e-erps-all-nodes.conf -configuration file for one-population neural field model.
All parameters are in SI units.
Time: 0.25 Deltat: 2.44140625e-4
Nodes: 4096
    Connection matrix:
From: 1 2
To 1: 1 2
To 2: 0 0
Population 1: Excitatory
Length: 0.5
Q: 10
Firing: Function: Sigmoid Theta: 0.01292 Sigma: 0.0038 Qmax: 340
 Dendrite 1: alpha: 83 beta: 769
 Dendrite 2: alpha: 83 beta: 769
Population 2: Stimulation
Length: 0.5
  Stimulus: Superimpose: 2
    Stimulus: Pulse - Onset: 0.03125 Node: 2000 Amplitude: 2
                      Width: 0.001953125 Frequency: 1 Pulses: 1
    Stimulus: Pulse - Onset: 0.06250 Node: 2097 Amplitude: −2
                      Width: 0.001953125 Frequency: 1 Pulses: 1
Propagator 1: Wave - Tau: 0 Range: 0.2 gamma: 30
Propagator 2: Map -
Coupling 1: Map -nu: 0
Coupling 2: Map -nu: 1e-4 32
Output: Node: All Start: 0 Interval: 9.765625e-4
Population: Dendrite:
Propagator: 1.phi
Coupling:
~~~

## Acknowledgments

The authors thank B. Fulcher, M. Prodanovic, J. A. Roberts and R. G. Townsend for useful discussions and feedback.

